# Discovery and characterization of a specific inhibitor of serine-threonine kinase cyclin dependent kinase-like 5 (CDKL5) demonstrates role in hippocampal CA1 physiology

**DOI:** 10.1101/2023.04.24.538049

**Authors:** Anna Castano, Margaux Silvestre, Carrow I. Wells, Jennifer L. Sanderson, Carla A. Ferrer, Han Wee Ong, Yi Liang, William Richardson, Josie A. Silvaroli, Frances M. Bashore, Jeffery L. Smith, Isabelle M. Genereux, Kelvin Dempster, David H. Drewry, Navjot S. Pabla, Alex N. Bullock, Tim A. Benke, Sila K. Ultanir, Alison D. Axtman

## Abstract

Pathological loss-of-function mutations in cyclin-dependent kinase-like 5 (*CDKL5*) cause CDKL5 deficiency disorder (CDD), a rare and severe neurodevelopmental disorder associated with severe and medically refractory early-life epilepsy, motor, cognitive, visual and autonomic disturbances in the absence of any structural brain pathology. Analysis of genetic variants in CDD have indicated that CDKL5 kinase function is central to disease pathology. *CDKL5* encodes a serine-threonine kinase with significant homology to GSK3β, which has also been linked to synaptic function. Further, *Cdkl5* knock-out rodents have increased GSK3β activity and often increased long-term potentiation (LTP). Thus, development of a specific CDKL5 inhibitor must be careful to exclude cross-talk with GSK3β activity. We synthesized and characterized specific, high-affinity inhibitors of CDKL5 that do not have detectable activity for GSK3β. These compounds are very soluble in water but blood-brain barrier penetration is low. In rat hippocampal brain slices, acute inhibition of CDKL5 selectively reduces post-synaptic function of AMPA-type glutamate receptors in a dose-dependent manner. Acute inhibition of CDKL5 reduces hippocampal LTP. These studies provide new tools and insights into the role of CDKL5 as a newly appreciated, key kinase necessary for synaptic plasticity. Comparisons to rodent knock-out studies suggest that compensatory changes have limited the understanding of the roles of CDKL5 in synaptic physiology, plasticity and human neuropathology.

## Introduction

Pathological loss-of-function mutations in cyclin-dependent kinase-like 5 (*CDKL5*)[1, 2] cause CDKL5 deficiency disorder (CDD, OMIM 300203, 300672), a rare (incidence 1:40,000– 60,000[3–5]), severe neurodevelopmental disorder associated with severe early-life epilepsy, motor, cognitive, visual and autonomic disturbances[2, 6–9, 10]. Epilepsy in CDD does not respond to *any* antiseizure medications[11]; a newly approved therapy provides modest improvements[12]. Hemizygous males and heterozygous females can be equally and severely affected[13, 14]. These severe symptoms are in absence of any structural brain pathology[15]. CDKL5 is a key mediator of synaptic and network development and physiology. Recent analysis of genetic variants in CDD have indicated that CDKL5 kinase function is central to disease pathology[13, 16]. In other words, loss of kinase function seems functionally equivalent to loss of the protein.

In rodent models, CDKL5 is expressed throughout the CNS primarily in neurons, increases post-natally and is stabilized at peak levels in adults[17–19], indicating roles during development and into adulthood. These features make it impossible to tease apart the precise role of altered CDKL5 function in mediating these symptoms in CDD. It remains unclear whether symptomatic abnormalities are due to chronic CDKL5 dysfunction, acute CDKL5 dysfunction during an earlier critical developmental time point, worsened by epilepsy or any combination. Adult rodent models of CDD are associated with abnormal behaviors, visual disturbances[20] and multiple abnormal signaling cascades[21–25]. CDKL5 is localized to the dendritic spines of excitatory synapses as well as the nucleus of neurons[17, 26, 27]. CDKL5 functions in the formation of excitatory synapses with partners that include PSD-95[28], NGL-1[29], Shootin1[30], and actin[17]. Previous work had suggested CDKL5 specific substrates including MeCP2, DNMT1, AMPH1, NGL-1 and HDAC4 but reliable antibodies have not been available[20]. Recent work has identified MAP1S and microtubule end binding protein 2 (EB2) as physiological CDKL5 substrates in brain [19]. A phosphospecific antibody for EB2-S222 has been used to report CDKL5 activity in mouse models and human iPSCs [19, 31, 32]. CDKL5 is a binding partner of both PSD-95 and gephyrin[33]; binding with PSD-95 is critical for spine development[21]. Studies have typically found that loss of CDKL5 leads to a global reduction in excitatory synapse numbers[29, 34], reduced PSD-95[35, 36] and synapsin[35] with loss of AMPA-type glutamate receptors (GluA2[37]) and increased NMDA-type glutamate receptors (GluN2B[38]). Inhibitory synapses appear to be unaffected[29], although inhibitory synaptic currents are affected in some CDKL5 mouse models[39]. Embryonic knock-out of *Cdkl5* rodents either enhanced long-term potentiation (LTP) [38, 40] or did not affect LTP [37, 40] in an age dependent fashion. Reports so far do not fully explain the presumed excitation/inhibition imbalance in epilepsy. Further, immature CDKL5 deficient mice do not have seizures or epilepsy[21, 41] and appear insensitive to pro-convulsants such as kainate [21]. This paradox highlights a big gap in our understanding of CDKL5 function: early developmental onset of medically resistant epilepsy in CDD contrasts with a complete lack in experimental models. The reasons for this mismatch are unclear and include possible developmental or other rodent-specific compensations, which typically involve transcriptional changes. Finally, the role of acute CDKL5 kinase dysfunction at any developmental time point is unknown. To address this, we sought to develop a sensitive and specific inhibitor of CDKL5.

*CDKL5* is an X-linked gene encoding a 115 kDa serine-threonine kinase and member of the CMGC family that includes cyclin dependent kinases (<u>C</u>DK), <u>M</u>AP-kinases, glycogen synthase kinases (<u>G</u>SK) and cyclin-dependent kinase-like (<u>C</u>DKL) [42]. It has limited structural homology to CDKL1, 2, 3 and 4 (http://mbv.broadinstitute.org/) but significant homology to GSK3β[43]. *CDKL1-4* mRNA are expressed at much lower levels than *CDKL5* in mouse and human brain (http://mouse.brain-map.org). Of these kinases, only GSK3β has been linked to brain synaptic function[44].

CDKL5 appears to be linked to GSK3β function. *Cdkl5* knock-out mice have increased GSK3β activity, as evidenced by *hypo-*phosphorylation of GSK3β at S-9 and decreased β-catenin[22]. β-catenin, which is phosphorylated and destabilized by active GSK3β, is often used, with S9-phospho-GSK3β, as a marker of GSK3β activity[45]. Inhibition of GSK3β in CDKL5 (young but not old) knock-out mice normalizes expression of β-catenin and S9-phospho-GSK3β[22, 23]. While upstream signaling is also affected (mTOR and AKT)[21], it is completely unclear how CDKL5 and GSK3β activities are linked in CDKL5 knock-out mice. GSK3β mediates a yin-yang interaction of LTP and LTD. Activation of GSK3β is required for the induction of GluN-LTD while inhibition occurs with the induction of LTP[44]. Thus, development of a specific CDKL5 inhibitor must be careful to exclude crosstalk with GSK3β activity.

Here, we synthesized and characterized specific, high-affinity inhibitors of CDKL5 that do not have detectable activity for GSK3β. These compounds are very soluble in water, but blood-brain barrier penetration is low. When applied directly to rat hippocampal brain slices, acute inhibition of CDKL5 selectively reduces post-synaptic function of AMPA-type glutamate receptors in a dose-dependent manner and inhibits a key form of synaptic plasticity, hippocampal LTP. These studies provide new insights into the role of CDKL5 in neuronal function and pathology.

## Methods

### Synthetic Schemes

Compound **B1** was prepared as a TFA salt with >95% purity (by HPLC) according to Scheme 1 (Supplementary Information) via a reductive coupling reaction in the presence of sodium borohydride and ethanol followed by amide coupling and deprotection. Compounds **B4** and **B12** were prepared as an HCl salt with >95% purity (by NMR) according to Scheme 2 (Supplementary Information). An amide coupling reaction in the presence of propylphosphonic anhydride (T3P) followed by nitro group reduction generated a common intermediate. Subsequent amide coupling reaction followed by deprotection yielded **B4** and **B12**. Full characterization of the intermediates and final products (Figure 2) are included as Supplementary Information.

**Figure 1.**
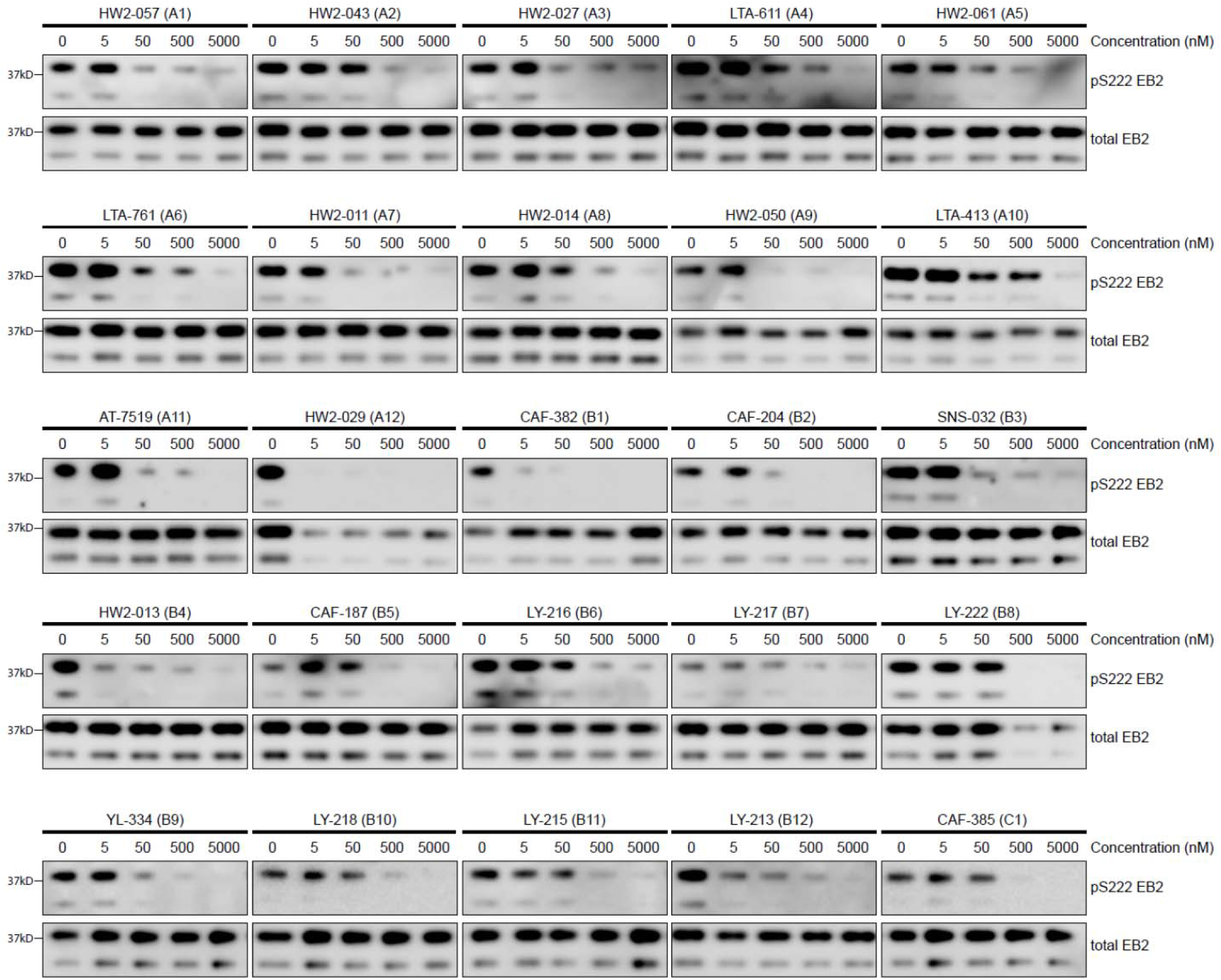
Screening of CDKL5 inhibitors in rat primary neurons using Western blotting. Western blots showing expression of total EB2 and levels of Ser222 EB2 phosphorylation in DIV14-16 rat primary neurons upon treatment of 1 hour with 5 nM, 50 nM, 500 nM and 5000 nM of selected CDKL5 inhibitors.

**Figure 2.**
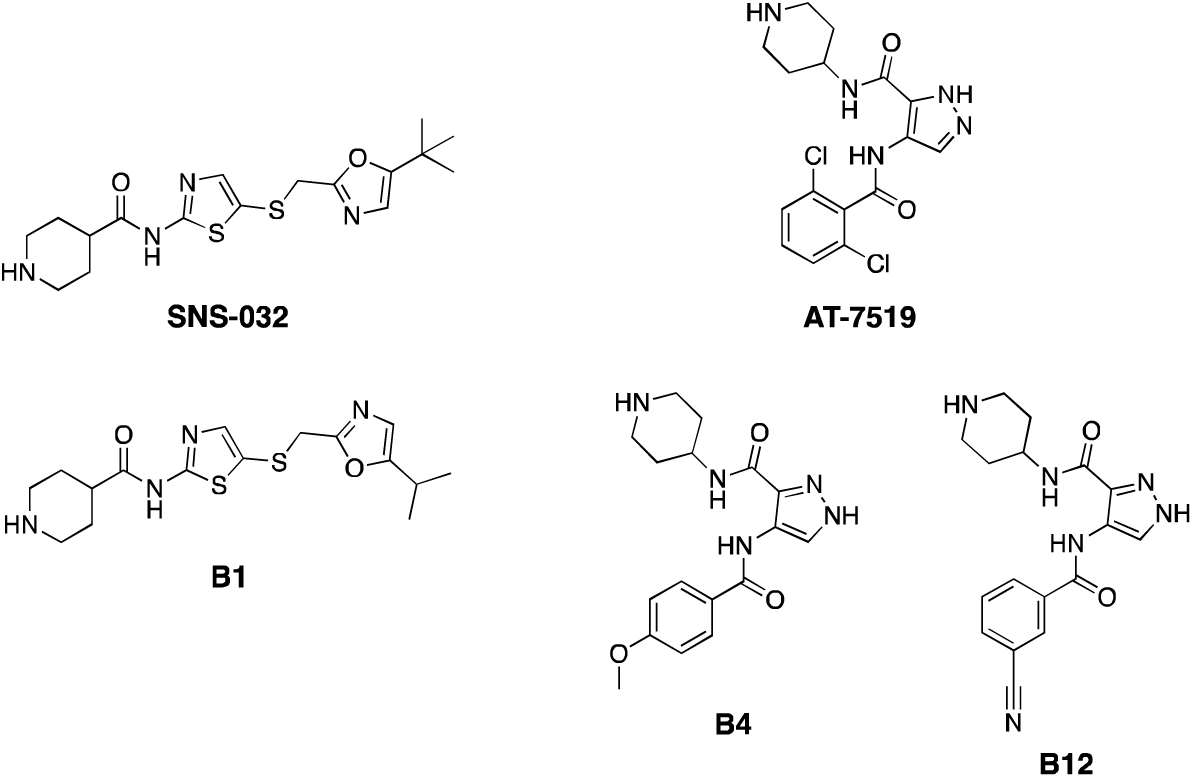
Structures of CDKL5 inhibitor leads and corresponding parent compounds.

### NanoBRET Assays

Human embryonic kidney (HEK293) cells obtained from ATCC (Manassas, VA, USA) were cultured in Dulbecco’s Modified Eagle’s medium (DMEM, Gibco) supplemented with 10% (v/v) fetal bovine serum (FBS, Corning). These cells were incubated at 37°C in 5% CO_2_ and passaged every 72 hours with trypsin (Gibco) so that they never reached confluency. Promega (Madison, WI, USA) kindly provided constructs for NanoBRET measurements of CDKL5 (NLuc-CDKL5), GSK31Z (NLuc-GSK31Z), and GSK3β (NLuc-GSK3β) included in Table S1 as well as Figures 3, S4, and S5. N-terminal NLuc orientations were used for all three kinases. NanoBRET assays were executed in dose–response (12-pt curves) format in HEK293 cells as previously reported [46–48]. Assays were executed as recommended by Promega, using 0.31 μM of tracer K11 for CDKL5, 0.13 μM of tracer K8 for GSK31Z, and 0.063 μM of tracer K8 for GSK3β.

**Figure 3.**
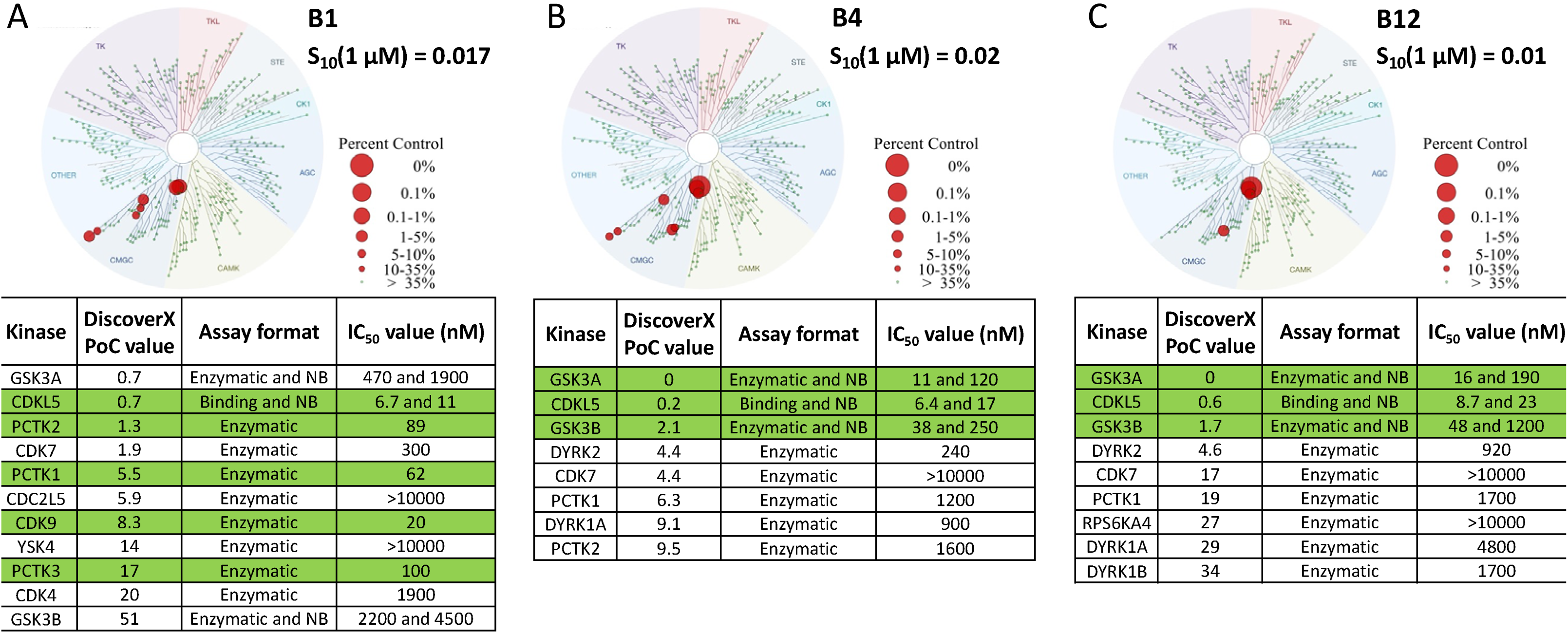
Kinome-wide selectivity data for CDKL5 lead compounds. Kinome tree diagrams illustrate the selectivity of these compounds when profiled against 403 wild-type (WT) human kinases at 1 µM at Eurofins DiscoverX in their *scan*MAX panel. The percent control legend shows red circles of different sizes corresponding with percent control value each kinase binds the small molecule in this large binding panel. Also included are selectivity scores (S_10_(1 µM)), which were calculated using the PoC values for WT human kinases in the *scan*MAX panel only. The S_10_ score is a way to express selectivity that corresponds with the percent of the kinases screened that bind with a PoC value <10. In the embedded tables kinases are listed by their gene names and ranked by their percent of control (PoC) value generated in the *scan*MAX panel. Rows colored green demonstrate enzymatic IC_50_ values within a 30-fold window of the CDKL5 binding IC_50_ value. Correlating assay format used to generate data in the last column of nested table is listed in the preceding column, with NB = NanoBRET.

### Analysis of Kinome-Wide Selectivity

The broad selectivity of compounds **B1**, **B4**, and **B12** at 1 µM was evaluated via the Eurofins DiscoverX Corporation (Fremont, CA, USA) *scan*MAX assay platform. In total, 403 wild-type (WT) human kinases were included in these analyses. Percent of control (PoC) values were generated, which allowed for the calculation of selectivity scores (S_10_(1 µM)) noted in Figure 3 [43]. WT human kinases in the *scan*MAX panel with PoC ≤20 (and GSK3β) for **B1**, with PoC <10 for **B4**, and PoC <35 for **B12** are also included in Figure 3. The kinome tree diagrams in Figure 3 were created based on WT kinases with PoC <10 for each compound.

### Biochemical Assays

A homogenous competition binding assay for CDKL5 was executed at Luceome Biotechnologies, LLC (Tuscon, AZ, USA) using their *KinaseSeeker* technology [49]. Briefly, this is a luminescence-based assay that relies on displacement of an active site dependent probe by an ATP-competitive test compound. Compounds **B1**, **B4**, and **B12** were evaluated in dose–response (12-pt curve) format in duplicate. Curves are included in Figure S4 and IC_50_ values generated are embedded in tables within Figure 3.

Radiometric enzymatic assays were executed at Eurofins at the K_m_ value for ATP to generate dose–response (9-pt) curves for all kinases listed in Figure 3 (except for CDKL5). A detailed protocol for these assays, which includes the protein constructs, substrate, and controls employed, is part of the Eurofins website: https://www.eurofinsdiscoveryservices.com.

### Kinetic Solubility

The kinetic solubility of compound **B1** was evaluated using an aliquot of a 10 mM DMSO stock solution dissolved in phosphate buffered saline solution (PBS) at pH 7.4. This analysis was done by Analiza, Inc (Cleveland, OH, USA) as described previously [50]. The reported solubility values in Table 1 have been corrected for background nitrogen present in DMSO and the media.

**Table 1.** Solubility and microsomal stability data for **B1**

| Compound | Kinetic solubility |  | Mouse liver microsomal stability (%) |  |  |
| --- | --- | --- | --- | --- | --- |
|  | µM | µg/mL | T = 0 min | T = 30 min | T = 30 min (-NADPH) |
| **B1** | 196.8<sup>[a]</sup> | 72.1<sup>[a]</sup> | 100 | 86.1 | 87.5 |

<sup>a</sup> Measured solubility is >75% dose concentration, actual solubility may be higher.

### Microsomal Stability

Mouse liver microsomal stability of compound **B1** was evaluated by Analiza, Inc (Cleveland, OH, USA) as described previously [48]. This data generated by Analiza is included in Table 1.

### CDKL Family Thermal Shift Assays

CDKL1, CDKL2, CDKL3, and CDKL5 proteins were produced using the construct boundaries previously used to generate 4AGU, 4AAA, 3ZDU, and 4BGQ, respectively. 4 µM of the kinase domain of CDKL1, CDKL2, CDKL3, or CDLK5 in 10 mM HEPES-NaOH pH 7.4 and 500 mM NaCl was incubated with **B1** (12.5, 25, or 50 µM) in the presence of 5× SyPRO orange dye (Invitrogen). Next, fluorescence was evaluated using a Real-Time PCR Mx3005p machine (Stratagene). A previously reported protocol was followed to run the Tm shift assays and assess melting temperatures[51]. Data is included in Figure S6.

### CDKL5 Enzymatic Assays

These assays were executed as previously described[52, 53]. Briefly, FLAG-tagged WT or kinase dead (KD, CDKL5 K42R) constructs of human CDKL5 were subcloned into a pT7CFE1-CHis plasmid (Thermo Fisher). A HeLa cell lysate-based Kit (1-Step Human Coupled IVT Kit—DNA, 88881, Life Technologies) enabled *in vitro* translation of these constructs. His Pur cobalt spin columns (Thermo Scientific) were used to purify the *in vitro*-translated proteins. To execute the *in vitro* kinase assays, myelin basic protein (Active Motif, 31314) was employed as a substrate for recombinant CDKL5. These two components were incubated in kinase buffer (Cell Signaling, 9802) supplemented with or without ATP (50 µM) for 30LJminutes at 30 °C, followed by kinase assays run using the ADP-Glo Kinase Assay kit (Promega). After the termination of kinase assay, NuPAGE LDS Sample Buffer (4X) was added to the reaction mixture and the samples were heated at 100°C for 10 minutes. The protein samples were run on Invitrogen Bis-tris gradient midi-gels and transferred to PVDF membrane for western blot analysis. Primary antibody used for western blot analysis was from Cell Signaling: FLAG (#14793) and was used at 1:1,000 dilution. Anti-Rabbit HRP-conjugated secondary antibody was from Jackson Immunoresearch (#111-035-144) and used at 1:5,000 dilution. Immunoblot signals were detected using SignalFire ECL Reagent (#6883) and X-ray films (Bioland #A03-02). Canon LiDE400 scanner was used for scanning the films. Precision Plus Protein Dual Color Standard prestained protein marker (Biorad) was used to estimate the molecular weight of sample proteins.

### Snapshot PK Study

The pharmacokinetics of compound **B1** was evaluated by Pharmaron (San Diego, CA, USA) following a single intraperitoneal administration to CD1 mice (two males). A dose of 2.29 mg/kg of the TFA salt of compound **B1** in NMP/Solutol/PEG-400/normal saline (v/v/v/v, 10:5:30:55) was prepared just before use. This dose was calibrated due to the compound being a salt form. Plasma was sampled 0.5-, 1-, 2-, and 4-hours post-dose. Bioanalytical assays were run using Prominence HPLC and AB Sciex Triple Quan 5500 LC/MS/MS instruments, and a HALO column (90A, C18, 2.7 µm, 2.1×50 mm). Snapshot PK results are included in Figure S10. PK parameters were estimated by non-compartmental model using WinNonlin 8.3 (Certara, Princeton, NJ, USA). The lower limit of quantification for plasma sampling was 1.53 ng/mL.

### Analysis of Brain/Plasma Concentration

The brain/plasma concentration of compound **B1** was evaluated by Pharmaron following a single intraperitoneal administration to CD1 mice (three males per dose). Two doses (2.29 mg/kg and 7.63 mg/kg) of the TFA salt of compound **B1** in NMP/Solutol/PEG-400/normal saline (v/v/v/v, 10:5:30:55) were prepared just before use. These doses were calibrated due to the compound being a salt form. Plasma and brain samples were collected 1-hour post-dose. Brain samples were homogenized at a ratio of 1:3 with PBS (W/V, 1:3) and then final brain concentrations corrected. Bioanalytical assays were run using Prominence HPLC and AB Sciex Triple Quan 5500 LC/MS/MS instruments, and a HALO column (90A, C18, 2.7 µm, 2.1×50 mm). Brain/plasma concentration results are included in Table S2. The lower limit of quantification for plasma sampling was 1.53 ng/mL and 0.763 ng/mL for brain.

### Animals-The Francis Crick Institute (TFCI)

Rat handling and housing was performed according to the regulations of the Animal (Scientific Procedures) Act 1986. Animal studies were approved by the Francis Crick Institute ethical committee and performed under U.K. Home Office project license (PPL P5E6B5A4B).

### Rat Neuron Primary Culture-TFCI

Primary cortical cultures were prepared from embryonic day (E) E18.5 embryos of Long Evans rats as described [54]. Pregnant females were culled using cervical dislocation, embryos were removed from the uterus and the brains were taken out. Cortices were dissected out, pooled from multiple animals, and washed three times with HBSS. An incubation with 0.25 % trypsin for 15 minutes at 37 °C was followed by four times washing with HBSS. Cells were dissociated and then counted using a hemocytometer. Neurons were plated on 12-well culture plates at a density of 300,000 cells per well. The wells were coated with 0.1 M borate buffer containing 60 µg/mL poly-D-lysine and 2.5 µg/mL laminin and placed in the incubator overnight. Neurons were plated with minimum essential medium (MEM) containing 10% fetal bovine serum (FBS), 0.5% dextrose, 0.11 mg/mL sodium pyruvate, 2 mM glutamine and penicillin/streptomycin. After 4 hours, cultures were transferred to neurobasal medium containing 1 mL of B27 (Gibco), 0.5 mM glutamax, 0.5 mM glutamine, 12.5 µM glutamate and penicillin/streptomycin. Primary neuronal cultures were kept at 37 °C and 5% CO_2_. Every 3-4 days, 20-30% of the maintenance media was refreshed. Between DIV14-16, neurons were treated with 5 nM, 50 nM, 500 nM and 5 µM of the different compounds for 1 hour. The compounds were added directly to the media and the plates were place at 37 °C for the time of the treatment. DMSO was added to the well for the control condition.

### Western Blotting-TFCI

After treatment, neuronal cultures were lysed in 300 µL of 1X sample buffer (Invitrogen) containing 0.1 M DTT. Lysates were sonicated briefly twice and denatured at 70 °C for 10 minutes. The samples were centrifuged at 13,300 rpm for 10 minutes and ran on NuPage 4-12% Bis-Tris polyacrylamide gels (Invitrogen). Proteins were transferred onto a Immobilon PVDF membrane (Millipore), which was then blocked in 4% milk in tris-buffered saline containing 0.1% Tween-20 (TBST) for 30 minutes. Primary antibodies were incubated at 4 °C overnight, and HRP-conjugated secondary antibodies at RT for 2 hours. The following primary antibodies were used: rabbit anti-CDKL5 (1:1,000; Atlas HPA002847), rabbit anti-pS222 EB2 (1:2,000; Covalab, from [19]), rat anti-EB2 (1:2,000; Abcam ab45767), mouse anti-tubulin (1:100,000; Sigma T9026), rabbit anti-β-catenin (1:1,000; Cell Signalling 9562) and rabbit anti phospho-β-catenin (Ser33/37 Thr41) (1:250, Cell Signalling 9561).The following secondary antibodies were used at a concentration of 1:10,000: HRP-conjugated anti-rabbit (Jackson 711-035-152), HRP-conjugated anti-mouse (Jackson 715-035-151) and HRP-conjugated anti-rat (Jackson 712-035-153). The membrane was developed using ECL reagent (Cytiva) and was visualized with an Amersham Imager 600 (GE Healthcare). Quantification of Western blots was manually performed using Image Studio Lite Software (version 5.2). EB2 phosphorylation was measured relative to total EB2 and catenin phosphorylation to total catenin. The other proteins were normalized to tubulin, if not indicated otherwise.

### Animals-University of Colorado School of Medicine (UC-SOM)

All studies conformed to the requirements of the National Institutes of Health *Guide for the Care and Use of Laboratory Rats* and were approved by the Institutional Animal Care and Use subcommittee of the University of Colorado Anschutz Medical Campus (protocol 00411). Timed-pregnant Sprague Dawley rats (Charles Rivers Labs, Wilmington, MA, USA) gave birth in-house. All rodents were housed in micro-isolator cages with water and chow available *ad libitum*.

### Hippocampal Slice Preparation and Electrophysiology-UC-SOM

As done previously [55–57], following rapid decapitation and removal of the rat brain at post-natal day (P) 20–30, sagittal hippocampal slices (400 µm) were made using a vibratome (Leica VT 1200, Buffalo Grove, IL) in ice-cold sucrose artificial cerebral spinal fluid (sACSF: 206 mM sucrose, 2.8 mM KCl, 1 mM CaCl_2_, 3 mM MgSO_4_, 1.25 mM NaH_2_PO_4_, 26 mM NaHCO_3_, 10 mM D-glucose and bubbled with 95%/5% O_2_/CO_2_). Following removal of CA3, slices were recovered in a submersion type chamber perfused with oxygenated artificial cerebral spinal fluid (ACSF: 124 mM NaCl, 3 mM KCl, 1 mM MgSO_4_, 2 mM CaCl_2_, 1.2 mM NaH_2_PO_4_, 26 mM NaHCO3, 10 mM D-glucose and bubbled with 95%/5% O_2_/CO_2_) at 28°C for at least 90 min and then submerged in a recording chamber perfused with recirculated ACSF. All electrophysiology was performed in the CA1 region at 28°C. Drugs were added to ACSF. Two twisted-tungsten bipolar stimulating electrodes were offset in the CA1 to stimulate one or two independent Schaffer collateral-commissural pathways using a constant current source (WPI, Sarasota, FL) with a fixed duration (20 µs), each at a rate of 0.033 Hz. Field excitatory post-synaptic potentials (fEPSPs) were recorded from the stratum radiatum (or stratum pyramidale region where indicated) region of CA1 using a borosilicate glass (WPI, Sarasota, Fl) microelectrode (pulled (Sutter, Novato, CA) to 6 to 9 MΩ when filled with ACSF), amplified 1000x (WPI, Sarasota, Fl and Warner, Hamden, CT), and digitized (National Instruments, Austin, Texas) at 20 kHz using winLTP-version 2.4[58] to follow fEPSP slope (averaged over 4 EPSPs), measured using 20% to 80% rise times, expressed as percent of baseline, during the course of an experiment. To be sure only “healthy” slices were included in our studies, responses had to meet several criteria: fiber volleys less than 1/3 of response amplitude and peak responses larger than 0.5 mV; responses and fiber volley must be stable (<5% drift). Following baseline stabilization of fEPSP slope at ∼50% of maximal slope for at least 30 min, measurements of synaptic physiology were performed. Stimulation was adjusted to make 4-6 measurements from ∼10% to -90% maximal slope to measure input-output (I/O) curves; recordings that deviated from linear slope were rejected. Paired-pulse ratios of slopes and peaks were measured at ∼50% of maximal slope. For measurements of NMDA-receptor dependent population spikes [59], Mg2+ was omitted from recording ACSF, the recording electrode was placed in stratum pyramidale, stimulation was adjusted to evoke a first population spike > 2 mV, and stabilization of the 2^nd^ NMDA-receptor dependent population spike (measured from the coast-line of the fEPSP) for greater than 30 minutes was allowed prior to measuring the effect of antagonists. Theta burst LTP was induced at test strength of two trains, separated by 20 seconds, of 10 trains of 4 stimuli at 100Hz, separated by 200 msec. Potentiation was statistically compared over the last 5 minutes.

For whole-cell, voltage-clamp electrophysiological recordings of excitatory post-synaptic currents (EPSCs), 350 μm horizontal hippocampal slices were prepared from rats at post-natal day (P) 20-30 similar to previously[60, 61]. After 15 min recovery at 34°C, slices were maintained at room temperature until recording in modified ACSF containing (in mM): 126 NaCl, 5 KCl, 2 CaCl_2_, 1.25 NaH_2_PO_4_, 1 MgSO_4_, 26 NaHCO_3_, 10 glucose and 2 N-acetyl cysteine. Slices were incubated in either CAF-382 (**B1**) (100 nM) or vehicle (water) in modified ACSF for at least 1h prior to recording. Using infrared–differential interference contrast microscopy for visualization, whole-cell recordings (Axopatch 200B amplifier and pClamp software (Molecular Devices)) from CA1 pyramidal neurons (series resistance between 3 and 6 MΩ) were performed at 29-30°C with an intracellular solution containing (in mM): 130 Cs gluconate, 1 CsCl, 1 MgS0_4_, 10 HEPES, 1 EGTA, 4 MgATP, 0.4 MgGTP, spermine (0.01) and 2 QX-314, pH 7.3. AMPA-type EPSCs, evoked using a bipolar tungsten-stimulating electrode as above, at a holding potential of −65 mV were pharmacologically isolated using picrotoxin (50 μM; Tocris). EPSC peaks were evoked every 20 seconds. After establishing whole-cell access, a 3 min baseline recording at −65 mV was followed by LTP induction using a pairing protocol: cells were depolarized to 0 mV for 90 sec and stimulated every 20 sec then followed by delivery of 1s x 100Hz high frequency stimulation. Evoked EPSC amplitudes were subsequently monitored at -65 mV. Potentiation was statistically compared over the last 5 minutes.

### Western Blotting-UC-SOM

Hippocampal slices were prepared and recovered as for electrophysiology with the addition of cuts to isolate CA1 from the remaining hippocampus; after recovery, the slices from a given rat were incubated in inhibitor or vehicle (water) in ACSF for 2 hours. Following this, slices were suspended in STE buffer, sonicated, boiled for 5 minutes and then frozen until further use. To minimize the effects of slice preparation[62] and to control for slice quality, only slices that came from a preparation that met electrophysiological criteria were used. All concentrations were quantified with BCA and then loaded in duplicate on 10% polyacrylamide gel and a five-point dilution series of naive rat hippocampal homogenate was included on each gel as a standard curve for quantification of immunoreactivity/ μg loaded protein[63]. Following transfer to PolyScreen PVDF transfer membrane (Genie, Idea Scientific Company, Minneapolis, MN, USA), blots were blocked in BSA or Carnation nonfat dry milk for 1 h and incubated either 1h at room temperature or overnight at 4°C with antibodies. The following primary antibodies were used: sheep anti-CDKL5 (1:2,000; University of Dundee, Anti-CDKL5(350-650)), rabbit anti-pS222 EB2 (1:1,000; Covalab, from [19]), rat anti-EB2 (1:1,000; Abcam ab45767). Blots were then subjected to three 10-min washings in Tris-buffered saline (140 mm NaCl, 20 mm Tris pH 7.6) plus 0.1% Tween 20 (TTBS), before being incubated with anti-sheep, anti-rat or anti-rabbit secondary antibody (1:5000) in 1% BSA or milk for 1 h at room temperature, followed by three additional 10-min washes preformed with TTBS. Immunodetection was accomplished using a chemiluminescent substrate kit (SuperSignal West Femto Maximum Sensitivity Substrate; Pierce) and the Alpha Innotech (Alpha Innotech, San Leandro, CA, USA) imaging system. Quantification was performed using AlphaEase software (Alpha Innotech, San Leandro, CA, USA) and Excel (Microsoft, Redmond, WA, USA). Immunoreactivity was reported as the density of sample bands relative to the standard curve. For phospho-proteins, ratios of immunoreactivity/μg to totals are reported without standardization. Only values falling within the standard curve generated from the dilution series included on each gel were incorporated into the final analysis. Some of the blots were then stripped (Restore PLUS Western Blot Stripping Buffer; Thermo Scientific, Rockford, IL) and reblotted if needed.

### Statistical Analysis-OSU

Data were analyzed via GraphPad Prism 9. Each assay was run in triplicate and mean values are graphed in Figure 4 with error bars showing standard deviation (SD). In Figure 4, statistical analysis was done using one-way ANOVA with Dunnett’s multiple comparisons test. Thresholds for significance (p-values) were set at ***p <0.0001 and non-significant statistics were not included.

**Figure 4.**
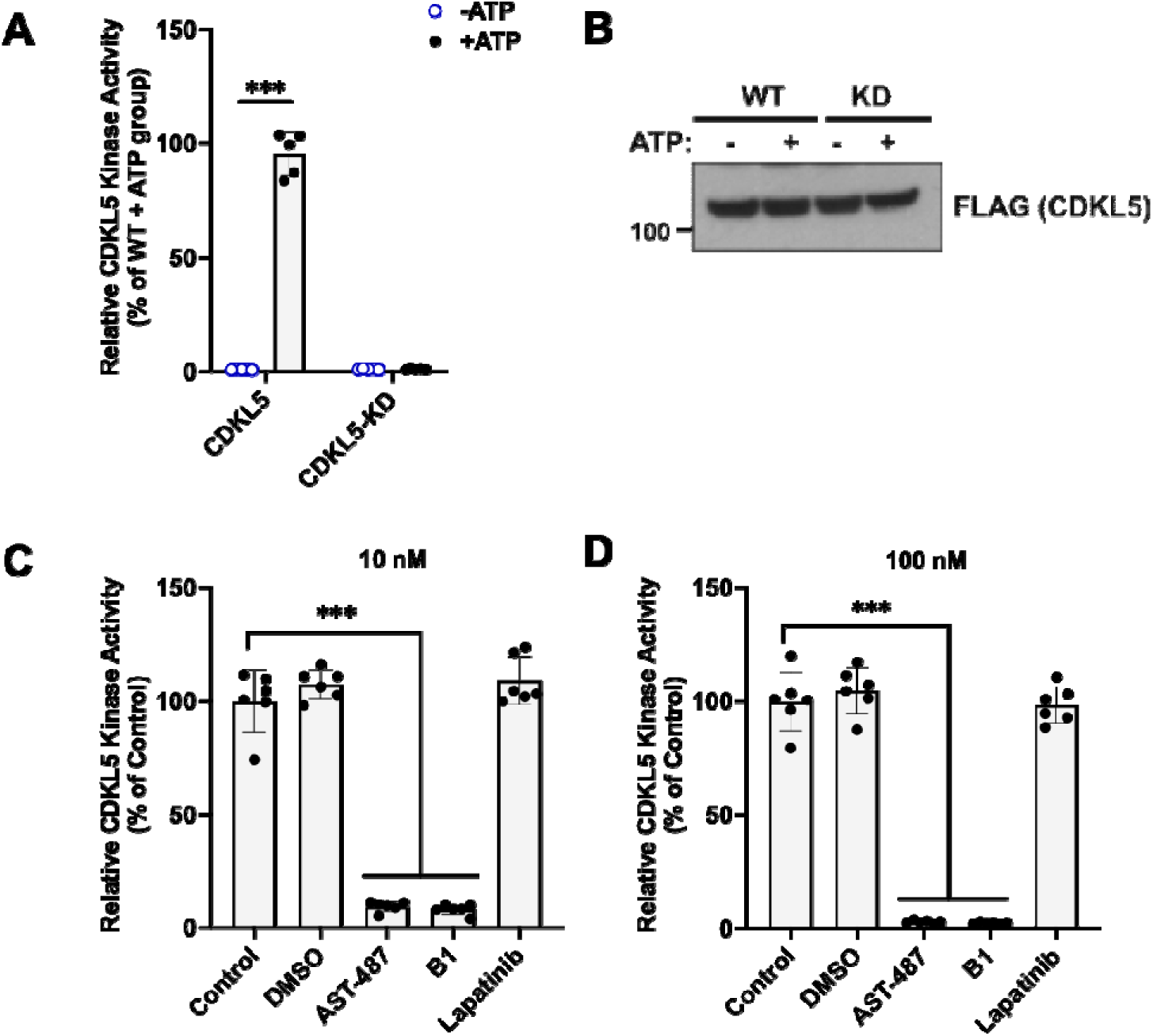
**B1** potently inhibits the kinase activity of CDKL5. B1 was simultaneously assayed with other CDKL5 antagonists[78]. (A) Representative graph from a CDKL5 kinase assay demonstrating that purified WT human CDKL5 retains kinase activity, while the kinase dead (KD, CDKL5 K42R) human protein is functionally inactive (n = 3). (B) Representative western blot illustrating that equal expression of WT and KD proteins was observed in the kinase assay experiments. (C and D) Kinase assays using purified WT human CDKL5 in the presence of 10 or 100 nM of the indicated compounds (n = 3). One-way ANOVA with Dunnett’s multiple comparison test used. *** = p <0.0001 and non-significant comparisons not shown. Control, DMSO, Positive (AST-487) and negative (Lapatinib) controls are as in[78].

### Statistical Analysis-TFCI

Data were analyzed using GraphPad Prism 9. Exact values of n and statistical methods are mentioned in the figure legends. Each concentration was compared to the control using a one-way ANOVA test. A p-value higher than 0.05 was not considered as statistically significant. Thresholds for significance were placed at *p ≤ 0.05, **p ≤ 0.01, ***p ≤ 0.001 and ****p≤ 0.0001. All errors bars in the figures are standard deviation (SD) where indicated. Non-significant statistics were not included.

### Statistical Analysis-UCSOM

Data are expressed as mean ± SEM with n = number of rats for a given treatment, unless otherwise stated. Data are plotted using Sigmaplot 12.5 (Systat, Chicago, IL, USA). Mann-Whitney Rank Sum, one-way, two-way and repeated measures (RM) ANOVA (Holm-Sidak post-hoc testing) and Student’s t-tests were used, where indicated and appropriate, for statistical comparisons for electrophysiological and biochemical using SigmaPlot 12.5 (Systat, Chicago, IL). Significance is reported at p < 0.05 unless otherwise stated. All errors bars in the figures are standard error (SE) where indicated. Non-significant statistics were not included.

### Data and materials availability

Data has been uploaded to Dryad (https://doi.org/10.5061/dryad.sn02v6x88). Newly synthesized compounds are available from the senior authors.

## Results

### Identification of CDKL5 inhibitors that lack GSK3β activity

SNS-032 and AT-7519 were initially identified as potent CDK inhibitors [43, 64]. We profiled our extensive library of SNS-032 and AT-7519 analogs using the CDKL5 NanoBRET assay. In parallel, the CDKL5 actives (IC_50_ <1 µM) were profiled using the GSK3β NanoBRET assay. Based on the data generated, a plate of 25 compounds was assembled. Criteria for inclusion on this plate included an IC_50_ <400 nM in the CDKL5 NanoBRET assay and >20-fold difference between the CDKL5 NanoBRET and the GSK3β NanoBRET IC_50_ values (Table S1).

### A targeted screen of CDKL5 inhibitors in neurons

A set of 20 CDKL5 inhibitors were selected based on their selectivity properties for CDKL5 versus GSK31Z/β for testing their inhibition of CDKL5 in rat primary cortical neuron cultures. We tested if we could detect a dose-dependent reduction of EB2 pSer222 following 1 hour incubation in culture media. We normalized pS222 EB2 with total EB2 levels to assess a specific reduction in phosphorylation as opposed to a loss of total protein levels. Almost all inhibitors showed a pSer222 EB2 reduction at 500 nM; most inhibitors showed a reduction at 50 nM (Figure 1). Three inhibitors caused a significant reduction in pSer222 EB2 at 5 nM without a change in total EB2 levels: CAF-382 (**B1**), HW2-013 (**B4**) and LY-213 (**B12**) (Figure 1). These represented different chemical parent backbones, so we decided to study these inhibitors further.

### Potency and selectivity analyses of lead compounds

As shown in Figure 2, **B1** is an SNS-032 analog, while **B4** and **B12** are analogs of AT-7519. SNS-032 and AT-7519 are published inhibitors of several members of the cyclin-dependent kinase (CDK) and cyclin-dependent kinase-like (CDKL) families [43]. Before using these compounds as tools for dissecting CDKL5 biology, we needed to first understand their kinome-wide selectivity. We analyzed **B1**, **B4**, and **B12** at 1 µM using the Eurofins DiscoverX *scan*MAX panel, which assesses binding to 403 wild-type (WT) human as well as several non-human and mutant kinases. Percent of control (PoC) values are generated for each kinase contained within the panel [43].

For **B1**, an analog of SNS-032, pan CDK inhibition was maintained (Figure 3A). All kinases with PoC ≤20 in the DiscoverX *scan*MAX panel plus GSK3β were profiled using orthogonal binding or enzymatic assays. While **B1** has a good selectivity score (S_10_(1 µM) = 0.017), several CDKs (CDK9, PCTK1/CDK16, PCTK2/CDK17, PCTK3/CDK18) are potently inhibited (IC_50_ ≤100 nM) (colored green in the nested table in Figure 3A). Weaker inhibition of CDK7 as well as both GSK31Z and GSK3β was noted. Curves corresponding to the affinity data for CDKL5 and GSK3 are included in Figures S4 and S5A, respectively. As the binding pockets share a high degree of similarly, the selectivity of **B1** within the CDKL family was further profiled via thermal shift assays (Figure S6). The highest change in melting temperature (ΔTm, >5 °C) was noted for CDKL5, while a modest ΔTm of 3–4 °C was observed for CDKL2. As a PoC value of 48 for CDKL2 was observed when **B1** was screened at 1 µM in the DiscoverX *scan*MAX panel, follow-up enzymatic studies were not executed for this kinase. We propose that only at higher concentrations (>10 µM) would inhibition of this kinase become significant.

Compounds **B4** and **B12**, which are analogs of AT-7519, also demonstrated good kinome-wide selectivity with S_10_(1 µM) scores of 0.02 and 0.01, respectively (Figure 3). Orthogonal profiling of **B4** confirmed high affinity binding to CDKL5, GSK31Z, and GSK3β (green in nested table in Figure 3B, curves in Figures S4 and S5B), with slightly weaker affinity for GSK31Z/β than CDKL5 evaluated via the respective cellular target engagement (NanoBRET) assays (Figures 3, S4, and S5). Kinases with PoC ≤10 in the DiscoverX *scan*MAX panel were profiled using orthogonal binding or enzymatic assays. An additional nine kinases demonstrated PoC <35. Orthogonal profiling of **B12** confirmed high affinity binding to CDKL5, GSK31Z, and GSK3β (green in nested table in Figure 3C, curves in Figures S4 and S5C). This compound is a more effective inhibitor of GSK31Z when compared to GSK3β. As was the case for **B4**, **B12** bound with greater affinity to CDKL5 than GSK31Z/β in the respective NanoBRET assays (Figures 3, S4, and S5C). Aside from GSK31Z/β, a large (>100-fold) window exists between CDKL5 and the next most potently inhibited kinase, DYRK2, when considering the enzymatic and/or binding data.

Binding and orthogonal assay results in combination with Western blots confirmed that in cells, **B1** is a CDKL5 and multi-CDK inhibitor with much weaker GSK31Z/β affinity (>1.8 µM) and inhibitory activity. Compounds **B4** and **B12** exhibited similar inhibition profiles and are potent inhibitors of CDKL5, GSK31Z, and GSK3β. Despite its potency, compound **B4** demonstrated sub-optimal selectivity when compared with **B1** and **B12**. This was determined based on the number of WT human kinases with PoC <35 in the DiscoverX *scan*MAX panel when profiled at 1 µM: **B1** = 12, **B4** = 17, **B12** = 9.

### In vitro kinase activity

We employed *in vitro* kinase assays to evaluate the impact of **B1** on CDKL5 activity (Figure 4). After preparing human WT and kinase dead (KD, CDKL5 K42R [52, 53]) recombinant proteins, equal amounts were employed in CDKL5 activity assays (Figure 4B). We observed that, in the presence of ATP (50 µM, Figure 4A), WT CDKL5 but not KD CDKL5 elicited kinase activity, as measured by turnover of ATP to ADP via a luminescent reagent (ADP-Glo assay). Next, the activity of untreated WT CDKL5 was compared with WT CDKL5 treated with vehicle (DMSO) or kinase inhibitors (10 or 100 nM). **B1**, AST-487, and lapatinib were the inhibitors chosen. AST-487 is a confirmed, but less selective inhibitor of CDKL5 (positive control) [65]. Lapatinib is a relatively selective kinase inhibitor that does not inhibit CDKL5 (negative control). The difference between DMSO, lapatinib, and untreated CDKL5 was not significant at 10 or 100 nM (Figure 4C, D). **B1** and AST-487, however, robustly inhibited CDKL5 in a dose-dependent manner (Figure 4C, D). **B1** was confirmed to be a potent inhibitor of CDKL5 kinase activity using this assay.

Kinase domains of GSK3β and CDKL5 have high similarity, to test the crosstalk of these inhibitors with GSK3β, we measured phospho-β-catenin (Ser33/37/Thr41), which is directly phosphorylated by GSK3β [19]. Phospho-β-catenin (Ser33/37/Thr41) is reduced upon exposure to GSK3β inhibitor CHIR 99021 while EB2 pSer222, is not affected by GSK3β inhibition (Figure S7). We measured pSer222 EB2 levels normalized to total EB2 at increasing concentrations of inhibitors in primary neurons and compared this to phospho-β-catenin normalized to total β−catenin (Figure 5). We found that CAF-382 (**B1**) showed a significant reduction in pSer222 EB2/total EB2 at 500 nM concentration (Figure 5A). There was a trend of pSer222 EB2/total EB2 reduction from 5 nM, which was not significant (Figure 5A). Phospho-β-catenin/ total β-catenin was not changed over the entire concentration range, indicating that CAF-382 (**B1)** does not inhibit GSK3β (Figure 5A). HW2-013 (**B4)** was effective in reducing EB2 pSer222 at 500 nM but also inhibited GSK3β significantly at 500 nM with a trend of inhibition from 5 nM, indicating that this inhibitor affects GSK3β (Figure 5B). Finally, LY-213 (**B12**) reduced pSer222 EB2 at 500 nM but also seemed to decrease β-catenin phosphorylation at this concentration. LY-213 (**B12**) showed comparable trends of inhibition for GSK3β and CDKL5 (Figure 5C). There were no significant differences in total CDKL5, EB2 or β-catenin levels during these treatments (Figure S8). We concluded that CAF-382 (**B1**) can inhibit CDKL5 activity without affecting GSK3 activity in neurons.

**Figure 5.**
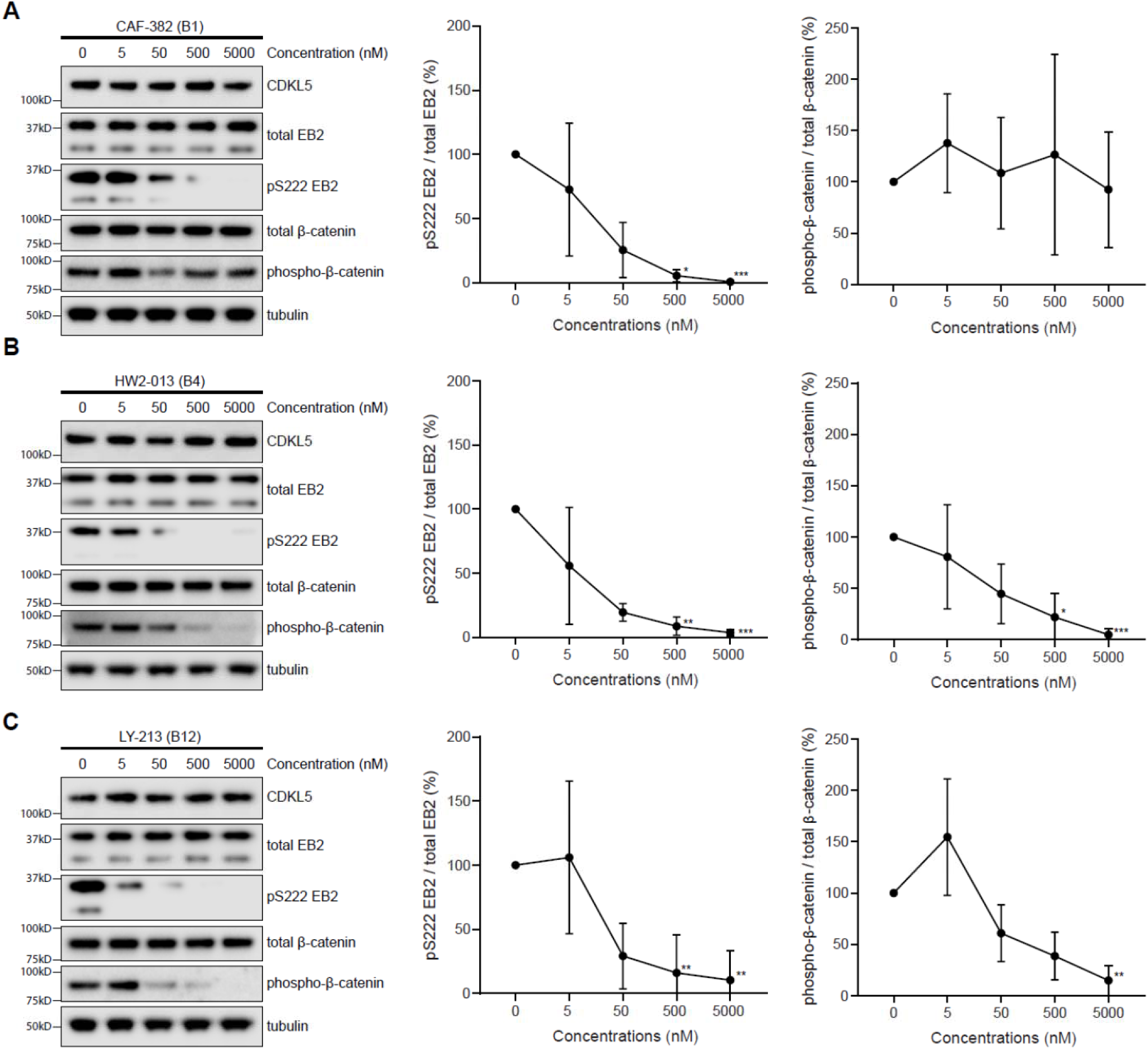
CAF-382 (**B1**), HW2-013 (**B4**), and LY-213 (**B12**) compounds reduce CDKL5 activity in rat primary neurons. HW2-013 (**B4**) and LY-213 (**B12**) also downregulate GSK3 activity. (A) Western blot and quantification showing expression of EB2 phosphorylation and β-catenin phosphorylation in DIV14-15 rat primary neurones after an hour treatment with different concentrations of CAF-382 (**B1**). (B) Western blot and quantification showing expression of EB2 phosphorylation and β-catenin phosphorylation in DIV14-15 rat primary neurons after an hour treatment with different concentrations of HW2-013 (**B4**). (C) Western blot and quantification showing expression of EB2 phosphorylation and β-catenin phosphorylation in DIV14-15 rat primary neurons after an hour treatment with different concentrations of LY-213 (**B12**). Each concentration was compared to the control using a Kruskal-Wallis test. n = 3 biological replicates with 2 repetitions. *p≤0.05. **p≤0.01. ***p≤0.001. Error bars are SD.

### Acute CDKL5 inhibition by CAF-382 (B1) in rat hippocampal slices

To determine if CAF-382 (**B1**) could inhibit CDKL5 activity within intact brain tissue, we prepared acute hippocampal slices from P20–30 rats and incubated them with CAF-382 (**B1**) for 2h as described. As for primary cultures, CAF-382 (**B1**) reduced EB2 phosphorylation in a dose dependent fashion (Figure 6). Similarly, SNS-032, the parent compound of CAF-382 (**B1**), also reduced EB2 phosphorylation in hippocampal slices (Figure S9).

**Figure 6.**
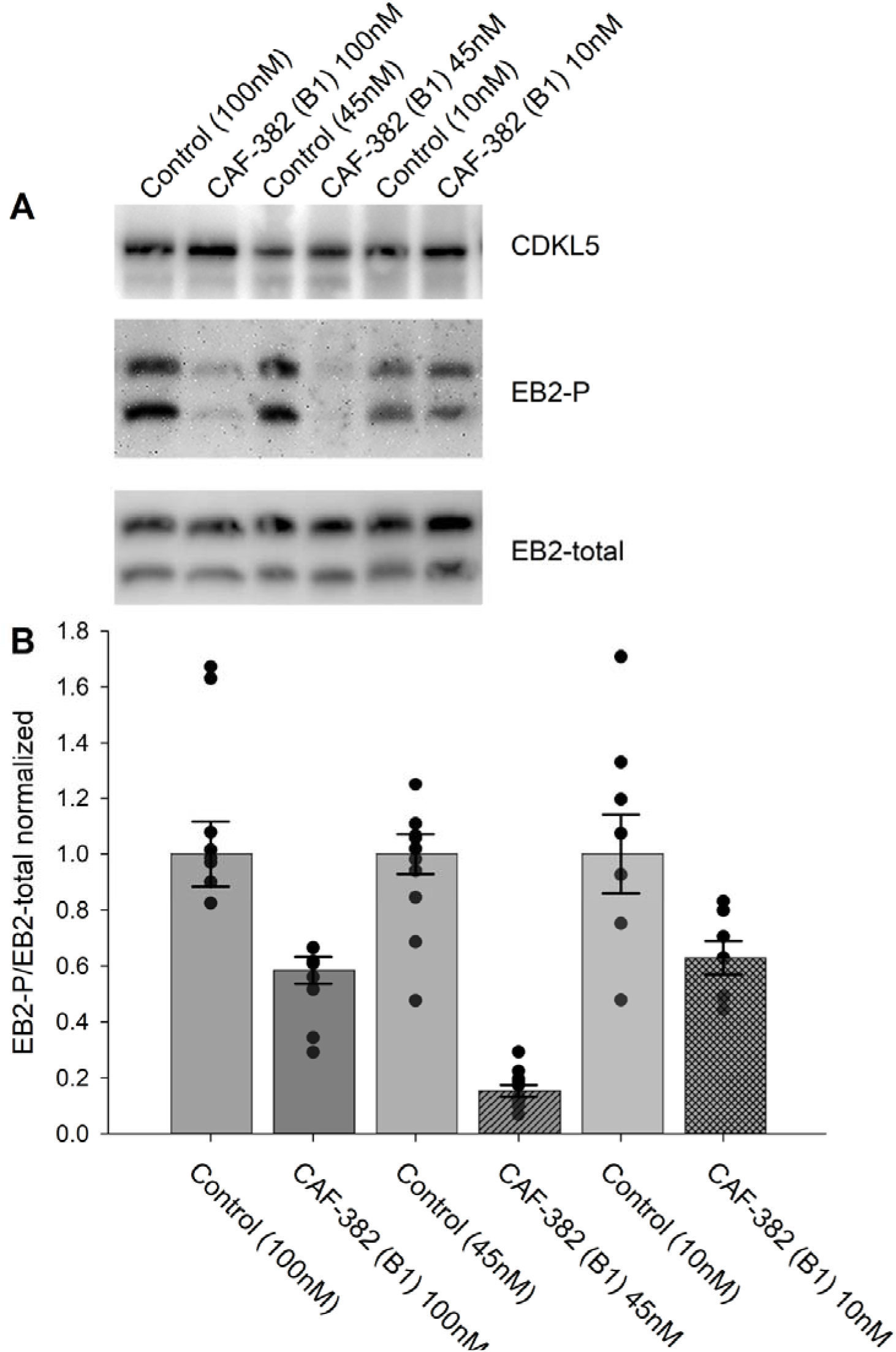
CAF-382 (**B1**) reduced phosphorylation of EB2 in CA1 hippocampal slices. (A) Example blots demonstrate no alterations in CDKL5 expression across treatments (stats in text) and suggest relative CAF-382 (**B1**) dose-dependent differences in EB2 phosphorylation. (B) Normalized quantification of EB2-phosphorylation (density of EB2-phosphorylation bands/ density of EB2-total bands). CAF 382 (**B1**) (control: 1±0.10 versus 100 nM: 0.49±0.02, n = 8, p = 0.002, RM-ANOVA) (control: 1±0.08 versus 45 nM: 0.16±0.03, n = 10, p < 0.001, RM-ANOVA) (control: 1± 0.14 versus 10 nM: 0.62± 0.11, n=7, p = 0.003, RM-ANOVA) reduced EB2 phosphorylation at all concentrations.

Previous studies found that genetic knock-down of *Cdkl5* in primary rodent brain cultures reduced glutamatergic excitatory synaptic transmission associated with loss of dendritic spines [29]. In this approach, the time-course of loss of CDKL5 function is expected to occur over several days. To clarify the more acute role of CDKL5 in regulating excitatory synaptic transmission, we applied CAF-382 (**B1**) (10–100 nM) to hippocampal slices while measuring fEPSPs in the CA1 hippocampus. While varying the stimulation strength applied to pre-synaptic Schaffer-collateral pathway inputs, we were able to measure the input-output responsiveness through measuring the input (fiber volley size) versus output (fEPSP slope). The slope of this relationship was reduced in a dose-response relationship by CAF-382 (**B1**) within 30 minutes of application (Figure 7A). In other words, for a similar input (fiber volley amplitude), the post-synaptic responsiveness (fEPSP slope) was reduced (Figure 7B). This suggests that the effect of acute CDKL5 inhibition by CAF-382 (**B1**) on glutamatergic synaptic transmission is primarily post-synaptic. Determination that amphiphysin-1, a pre-synaptic protein, is phosphorylated by CDKL5 [66] suggested a role of CDKL5 in presynaptic function. However, *Cdkl5* knock-out studies showed that amphiphysin-1 was not phosphorylated by CDKL5 *in vivo*, suggesting an alternative amphiphysin-1-independent mechanism is responsible for observed presynaptic changes [Kontaxi et al Mike Cousin, biorxiv]. To investigate the presynaptic role of CDKL5 kinase function, the paired-pulse ratio across several paired-pulse intervals was measured (Figure 8). CAF-382 (**B1**) (100 nM x 1 hour) had no effect on the paired-pulse ratio, confirming the acute effects of CAF-382 (**B1**) are restricted to the post-synapse. fEPSP slopes are mediated by AMPA-type glutamate receptors with NMDA-type glutamate receptors largely silent under these conditions. In order to visualize the impact of CAF-382 (**B1**) on NMDA-type glutamate receptor mediated responses, extracellular Mg^2+^ was removed from the recording medium [59]. Under these conditions, a NMDA-receptor mediated field potential (population spike 2) becomes apparent; this was not affected by CAF-382 (**B1**) (Figure 9) but was completely blocked by the NMDA-type glutamate receptor antagonist D-APV (50 μM). This result suggests that the acute effect of CAF-382 (**B1**) mediated CDKL5 inhibition is to selectively reduce AMPA-type glutamate receptor-mediated responses post-synaptically.

**Figure 7.**
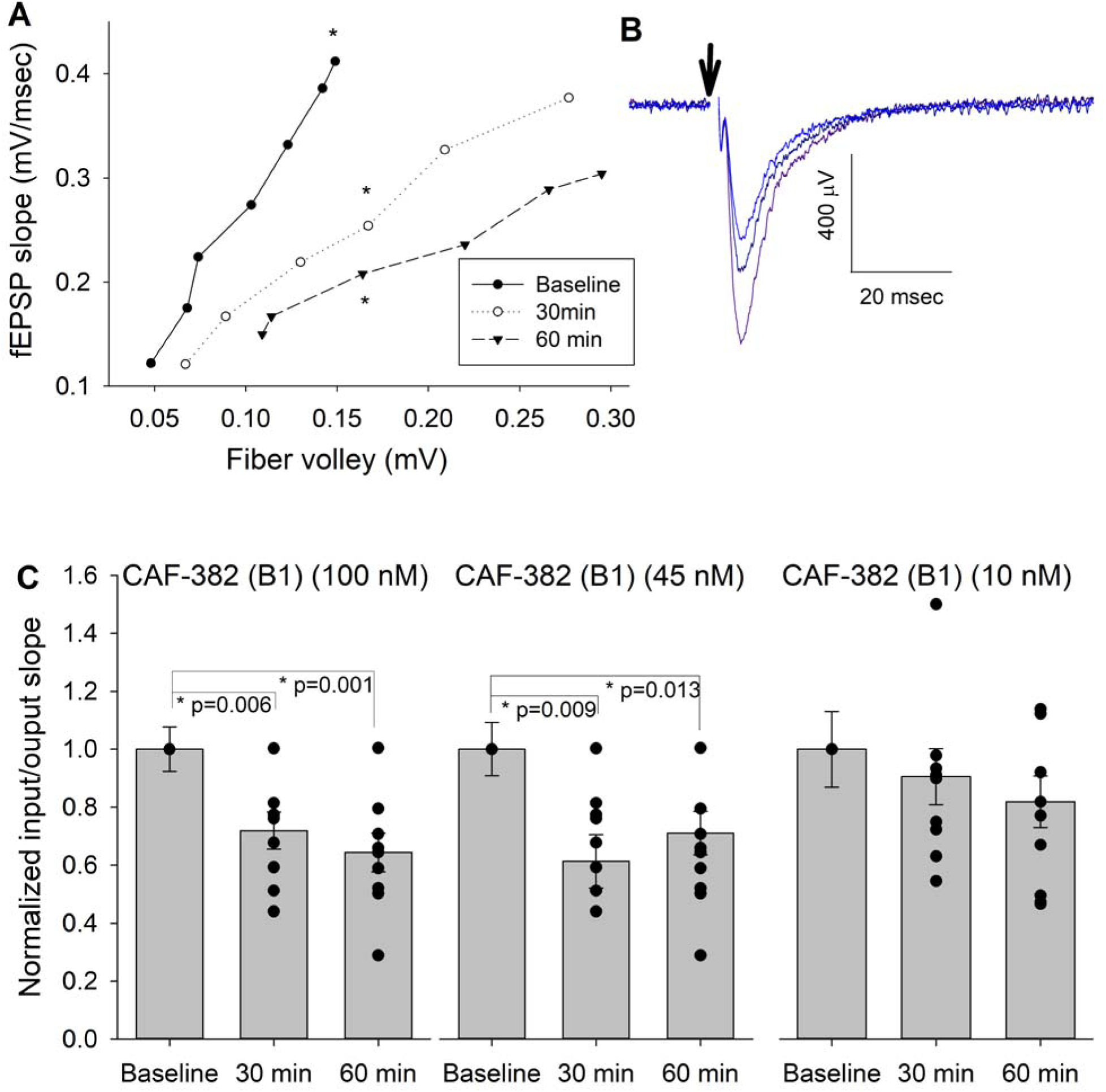
CAF-382 (**B1**) reduced post-synaptic fEPSP responsiveness, as measured by input-output curves. (A) Sample input (fiber volley) / output (fEPSP slope) (I/O) curves at baseline, 30 minutes and 60 minutes after CAF-382 (**B1**) (100nM) in a hippocampal slice. Slopes of I/O curves were obtained at baseline, and 30 minutes and 60 minutes after treatment with CAF-382 (**B1**). Asterisks indicate sample traces (B) from treated slice; stimulus artifact (arrow) has been removed for clarity. Initial negative deflection after the artifact is the fiber volley, followed by the fEPSP. (C) RM-ANOVA of I/O slopes of CAF-382 (**B1**) (100 nM and 45 nM) were significantly decreased after 30 minutes (100 nM: 1.91±0.23, n = 9, p = 0.006; 45 nM: 1.58±0.24, n=10, p = 0.013) and 60 minutes (100 nM: 2.00±0.30, n = 9, p = 0.001; 45 nM: 1.57±0.22, n=10, p = 0.009) compared to baseline (100 nM: 3.24±0.52, n = 9; 45 nM: 2.29±0.23, n=10). Baseline I/O slope was used to normalize each recording to allow comparisons across all treatments. CAF-382 (**B1**) (10 nM) did not alter I/O slopes after 30 or 60 minutes (RM-ANOVA, n = 9, p = 0.07). Error bars are SE.

**Figure 8.**
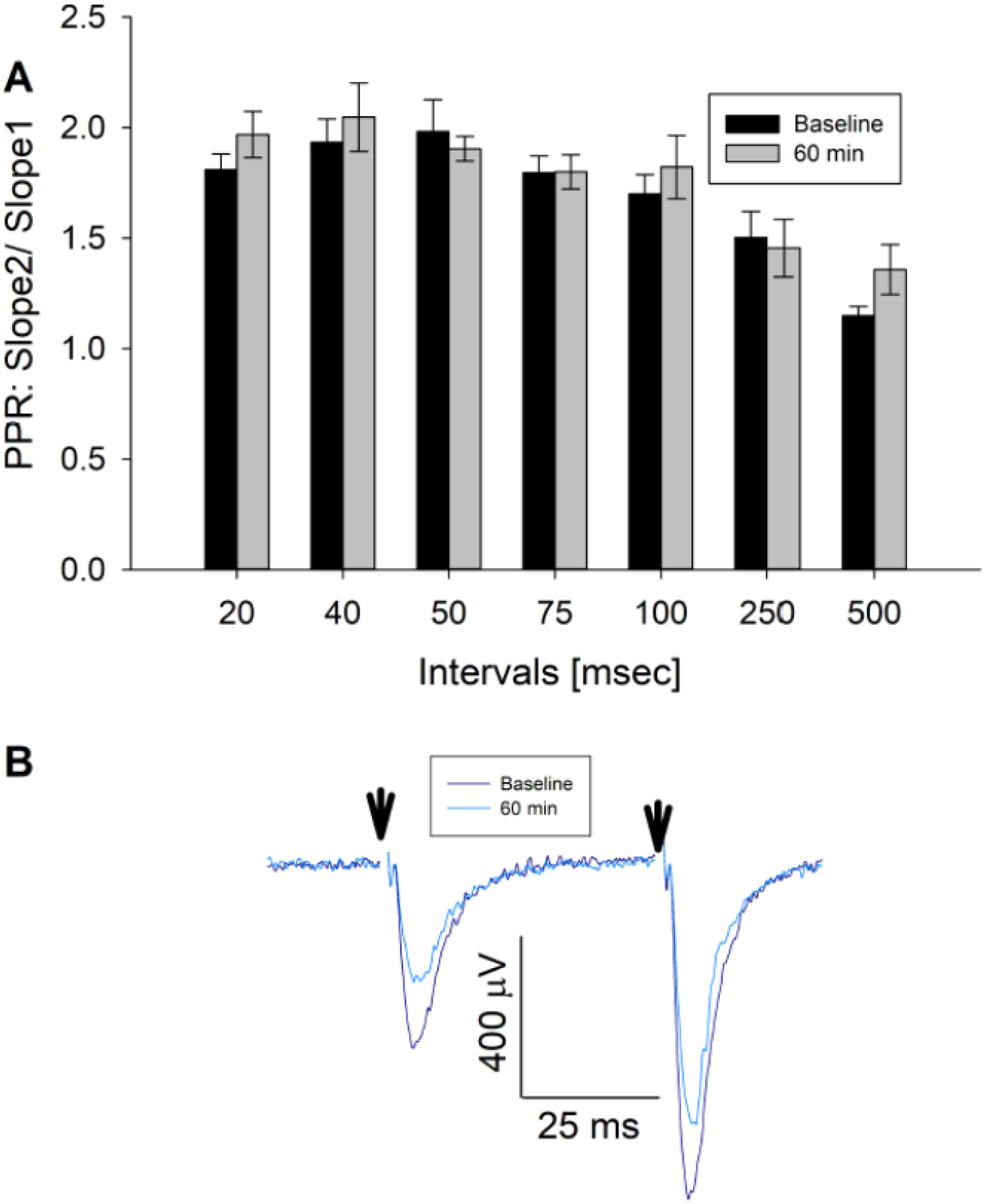
CAF-382 (**B1**) did not alter presynaptic release as measured by paired-pulse ratio. (A) Paired-pulse ratios (PPR) of fEPSPs (Slope of fEPSP2/Slope of fEPSP1), reflective of pre-synaptic release, were unaltered across a range of stimulus intervals (RM-ANOVA, n=7) by CAF-382 (**B1**) (100nM) for 60 minutes compared to baseline. (B) Sample trace for 50 ms interval. Reduction of initial fEPSP slopes were consistent with Figure 7. Stimulus artifacts (arrows) have been removed for clarity. Error bars are SE.

**Figure 9.**
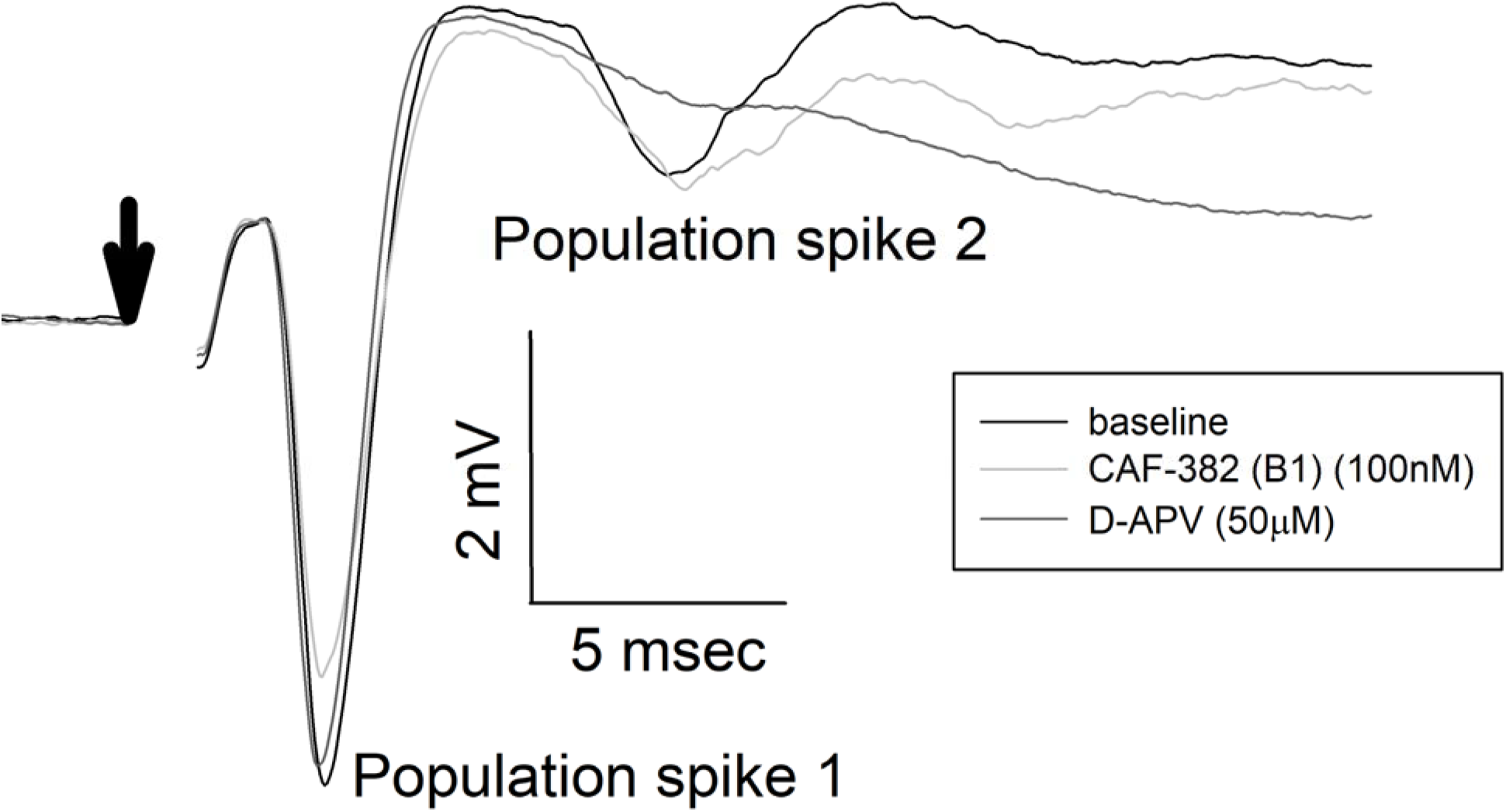
CAF-382 (**B1**) did not alter NMDAR-mediated component of fEPSPs. Maximal fEPSPs recorded in stratum pyramidale of CA1 with reduced extracellular Mg^2+^ demonstrate multiple population spikes; previous studies have demonstrated that secondary population spikes in these conditions are NMDAR-dependent[59]. CAF-382 (**B1**) (100 nM) did not alter population spike 1 (control -4.057±0.584mV, CAF-382 (**B1**) 3.929±0.736mV, n=10, p=0.742, RM-ANOVA) or NMDAR-mediated population spike 2 (control -2.093±0.568mV, CAF-382 (**B1**) - 1.656±0.4442, n=10, p=0.867, RM-ANOVA). For comparison, the selective NMDAR antagonist D-APV (50 μM) completely blocked population spike 2, as previously reported [59]. Stimulation artifact has been removed (arrow) for clarity.

A previous study found that germ-line knock-out of *Cdkl5* in the mouse enhanced long-term potentiation (LTP) at 7–13 week old (wo) hippocampal Schaffer Collateral to CA1 synapses without alteration of the input-output relationship [38]. In another study, germ-line knock-out of *Cdkl5* in a mouse at 4–5 wo did not affect stable LTP at this synapse [37]. In a rat *Cdkl5* knock-out, LTP at this synapse was selectively increased at 3–4 wo but not at later ages [40]. To clarify the more acute role of CDKL5 in regulating LTP at this synapse, we incubated 3– 4 wo hippocampal slices with CAF-382 (**B1)** (100 nM) for at least 30 minutes and, following stable baseline measurements of fEPSP slope, induced LTP via theta burst to Schaffer Collateral to CA1 synapses in the continuous presence of CAF-382 (**B1)**. Compared to interleaved recordings from control slices, CAF-382 (**B1)** significantly reduced LTP at 60 minutes post theta-burst (Figure 10). To investigate this further, LTP at these synapses of isolated AMPA-type glutamate receptors via a pairing protocol was measured using whole-cell patch-clamp. In this case, hippocampal slices were incubated with CAF-382 (**B1)** (100 nM) for only one hour prior to, but not during, recording. Compared to interleaved recordings from control slices, CAF-382 (**B1)** significantly reduced LTP at 30 minutes post pairing (Figure 11). The induction and expression of LTP includes the contributions of pre-synaptic and post-synaptic mechanisms at both excitatory and inhibitory synapses[67, 68]. Since we do not see a presynaptic effect of CAF-382 (**B1)** (Figure 8), and the approach (Fig. 11) removed the contribution of inhibitory synapses, this supports the direct role of CDKL5 in the post-synaptic mediation of LTP at this synapse at this age.

**Figure 10.**
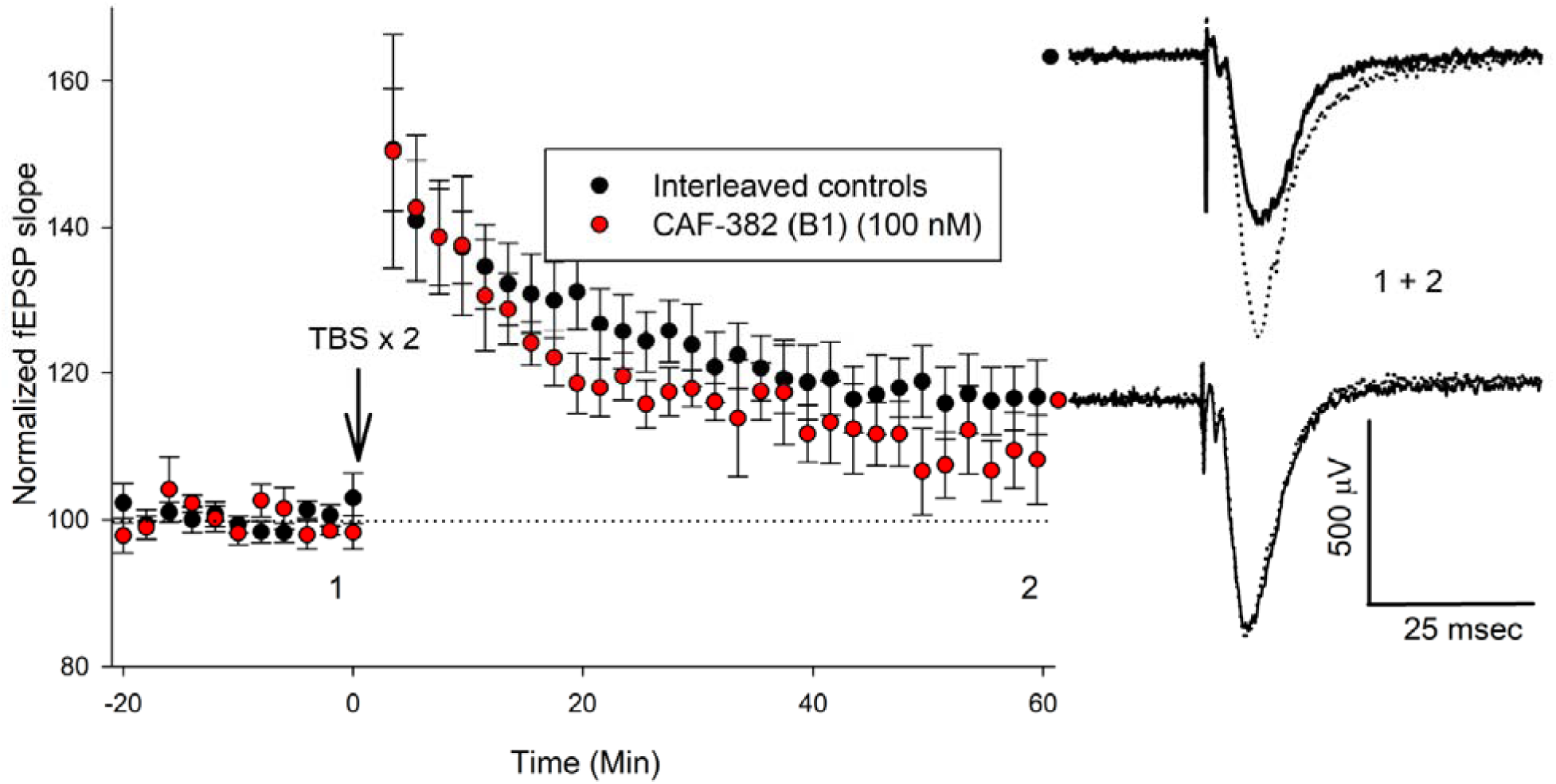
CAF-382 (**B1**) acutely reduces the expression of LTP in CA1 hippocampus. Hippocampal slices were initially incubated in CAF-382 (**B1**) for at least 30 minutes and then continuously perfused with CAF-382 (**B1**) (100 nM). fEPSP slope was normalized to initial baseline, expressed as 100% (dotted line); mean±SEM shown. Following stable baseline of fEPSP slope (> 20 minutes), LTP was induced by two theta-burst (TBS) trains. Compared to interleaved control slices, CAF-382 (**B1**) significantly reduced LTP as measured by fEPSP slope at 56-60 minutes post theta-burst (control: 115.6±2.2 n = 13 slices, 12 rats; CAF-382 (**B1**): 106.4±3.5, n = 5 slices, 4 rats; p = 0.029, 2-way ANOVA, Holm-Sidak post-hoc). Sample traces for control (black filled circle) and CAF-382 (**B1**) (red filled circle) before (1, solid trace) and after LTP (2, dotted trace) are shown to the right. Error bars are SE.

**Figure 11.**
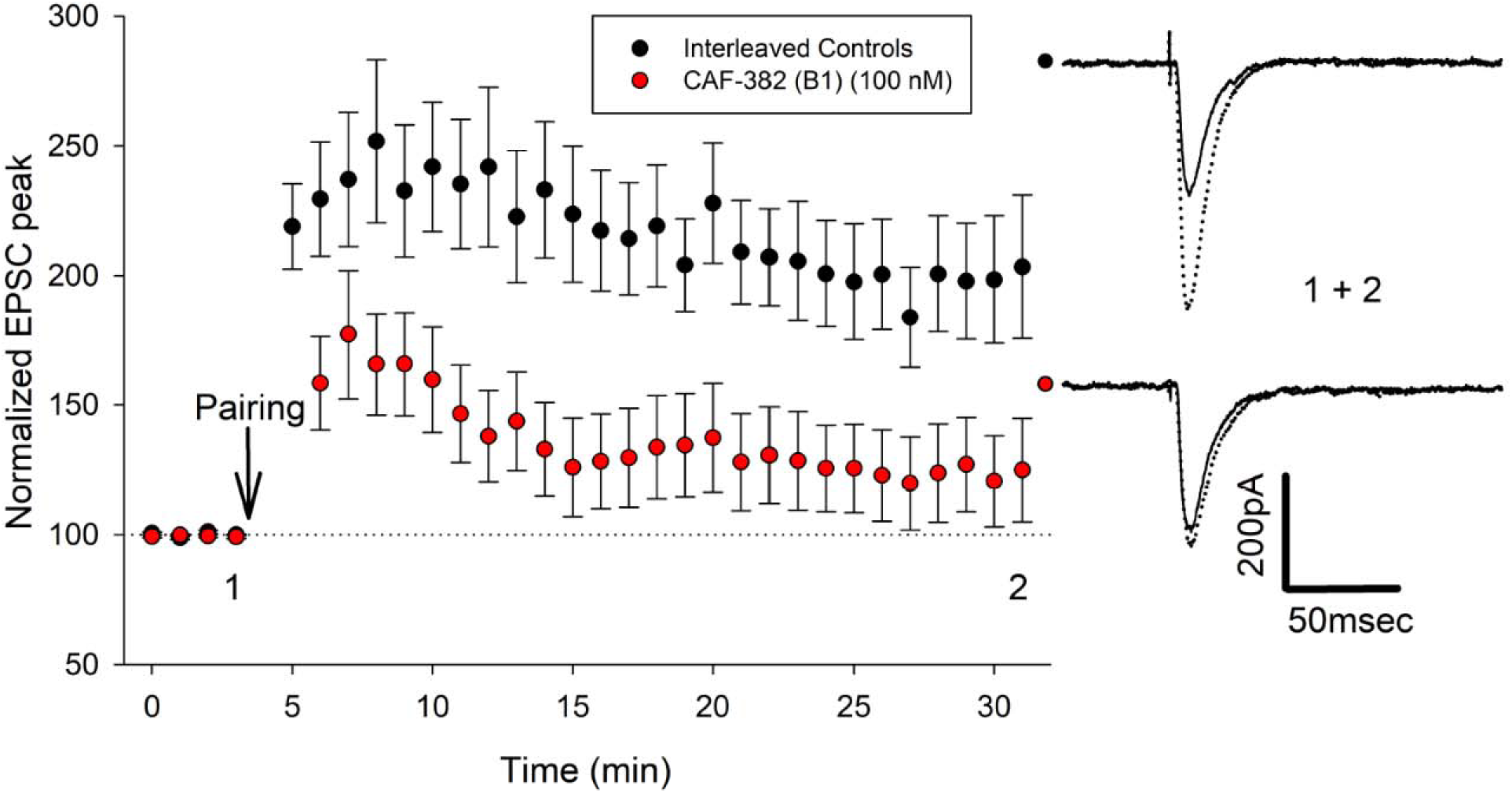
CAF-382 (**B1**) acutely reduces the expression of LTP mediated by AMPA-type glutamate receptors in CA1 hippocampus. Hippocampal slices were initially incubated in CAF-382 (**B1**) (100 nM) for at least 60 minutes prior to recording. Peak negative current of AMPA-type glutamate receptor mediated synaptic responses were normalized to initial baseline post break-in (time = 0), expressed as 100% (dotted line); mean±SEM shown. Following baseline, LTP was induced by a pairing protocol. Compared to interleaved control slices, CAF-382 (**B1**) significantly reduced LTP at 27-31 minutes post break-in (control: 197.5±8.4 n = 14 slices, 14 rats; CAF-382 (**B1**): 123.14±9.0, n = 12-13 slices, 13 rats; p < 0.001, 2-way ANOVA, Holm-Sidak post-hoc). Sample traces for control (black filled circle) and CAF-382 (**B1**, red filled circle) before (1, solid trace) and after LTP (2, dotted trace) are shown to the right. Error bars are SE.

### Evaluation of suitability of CDKL5 lead for in vivo studies

As our studies were all executed *in vitro*, the suitability of **B1** for use *in vivo* was unknown. To gauge whether **B1** could be dosed in animals using a water-based formulation, we first analyzed its aqueous kinetic solubility (Table 1). **B1** demonstrated good solubility (196 µM), supporting that it readily dissolves in aqueous buffer without observed precipitation. Mouse liver microsomal stability data was also collected (Table 1). **B1** proved to be very tolerant to microsomal exposure and demonstrated excellent stability (>85%) after 30 minutes. Since compound **B1** was found to have the required aqueous solubility and mouse liver microsomal stability, it was submitted for *in vivo* characterization. Pharmacokinetic analysis and brain exposure of mice following intraperitoneal administration with **B1** were next examined. Plasma was sampled 0.5-, 1-, 2-, and 4-hours post-dose with 2.29 mg/kg of a salt form of **B1** (Figure S10). No abnormal clinical symptoms were observed in dosed animals. The half-life of **B1** was found to be <1h, with the highest concentration of drug in the plasma found 0.5 h after dosing (Cmax = 508 ng/mL) and total system exposure (AUCinf) of 646 h*ng/mL (Figure S10). The same dose (2.29 mg/kg) as well as a 7.63 mg/kg dose were used to analyze the brain penetration of **B1** in mice 1h after administration. As shown in Table S2, brain and plasma sampling indicated that brain penetration of **B1** was generally low.

## Discussion

CDD is a rare, devastating, genetically mediated neurological disorder with symptomatic onset in very early infancy. Though there are newly approved therapies that provide some relief for epilepsy [12], the additional features that are impactful to caregivers (Jeff Neul, personal communication) of motor, cognitive, visual and autonomic disturbances [2, 6–9, 10] are not currently addressed. CDKL5 protein [69] and *CDKL5* gene [70] replacement approaches are advancing through pre-clinical studies to potential first-in-human trials (https://www.ultragenyx.com/our-research/pipeline/ux055-for-cdd/). While CDKL5 is a key mediator of synaptic and network development and physiology, the precise role of this kinase as explored through genetic manipulations in rodent models remains unclear. This knowledge gap prevents a full understanding of the role of CDKL5 across development and has implications for determining and interpreting when protein and gene replacement strategies may be most efficacious. As CDKL5 kinase function is central to disease pathology [13, 16], our development of a specific CDKL5 kinase inhibitor aims to address this gap.

Following identification of existing inhibitors [64], detailed analysis of the activity of analogs of these inhibitors in a range of assays identified three compounds with significant selectivity over GSK3β, a key requirement due to structural homology [43] and linkage of GSK3β to both synaptic physiology [44] and CDKL5-related signaling [22, 23]. The lead compound, **CAF-382 (B1**), demonstrated potent activity in multiple assays, including target-based *in vitro* biochemical assays and functional assays utilizing both rodent neuronal cultures and brain slices. The results across all assays are consistent, and align with the hypothesis that compound **B1** inhibits CDKL5 and blocks phosphorylation of a known CDKL5 substrate, EB2 [19, 31, 32]. Several cyclin dependent kinases (CDKs) that are potently (IC_50_ ≤100 nM) inhibited (CDK9 [71], PCTK1/CDK16 [72], PCTK2/CDK17 [73], and PCTK3/CDK18 [74]) by **B1** are expressed in the brain (http://mouse.brain-map.org). However, none of these have been demonstrated to have synaptic activity or localization; each are largely involved in replication and nuclear functions. At issue, is whether observations of CDKL5 activity, either through genetic manipulations or interference with the kinase, are specific and not potentially subsumed by the activity of other kinases. Given the reduction of phosphorylation of the specific substrate, EB2, this is less likely here.

The biochemical activity of **CAF-382** (**B1**) against CDKL5 is <5 nM, based on *in vitro* activity studies (Figure 4), and for studies investigating cellular target engagement as well as activity in neuronal cultures and brain slices at the specific substrate, EB2, and synaptic physiology we observe a 10-fold shift in potency (∼50 nM). A loss in potency, albeit modest, when moving from a biochemical to cell-based assay is commonly observed, mostly due to suboptimal cell penetrance of the small molecule, and has been observed for several orthogonal chemical series of kinase inhibitors[47, 48, 50]. As the *in vitro* assay eliminates the contribution of phosphatases (and the phosphatases specific for EB2 are currently unknown), the apparent mismatch could also be due to residual presence of phosphatase-dependent and previously phosphorylated EB2 balanced with active phosphorylation by CDKL5. Nevertheless, CDKL5 kinase function is clearly necessary for the induction and/or expression long-term potentiation, adding to the very short list of kinases required for this essential process linked to learning and memory[67, 68, 75] and finally clarifying previous genetic studies. The increased LTP seen in genetic studies suggests that observations of increased LTP in these models is compensatory and not directly due to loss of CDKL5. One interpretation of those prior studies is that CDKL5 provided a limiting “brake” on LTP, with genetic loss resulting in more LTP which is in contradistinction to our findings. The role of CDKL5 at inhibitory synapses was not specifically examined. Nevertheless, synaptic function at the developmental age studied (P20–30) requires CDKL5 activity for synaptic plasticity, suggesting that strategies that boost function at this age-equivalent in humans may be clinically important. Selective stability and plasticity dependent insertion of AMPA-type glutamate receptors at CA1 hippocampal synapses likely require a balance between active CDKL5 kinase and phosphatases. Further studies are needed to determine this balance more specifically and continue to clarify CDKL5 function at pre-synapses[76].

While no clinical abnormalities were observed with *in vivo* administration of CAF-382 (**B1**), this is likely due to low brain penetration. Due to low brain penetration of CAF-382 (**B1**), strategies that optimize brain penetration or different administration routes could be considered in future studies investigating the role of CDKL5 *in vivo*. These studies, behavioral and perhaps with electrical assessments of seizure activity, may clarify the apparent mismatch of rodent knock-out with clinical observations. Using CDKL5 inhibitors *in vivo* across developmental stages can help ascertain possible therapeutic windows for CDD. More broadly, CDKL5 inhibition is beneficial for cell survival upon ischemia or nephrotoxin induced kidney injury[53, 77], suggesting that the inhibitors we characterized here may prove useful to evaluate the promise of CDKL5 inhibition in other disease models.

## Supporting information

reviewer comments

## Funding

NIH-NINDS NS112770 (all), Ponzio Family Chair in Neurology Research (AC, TAB), The Structural Genomics Consortium (SGC) is a registered charity (number 1097737) that receives funds from Bayer AG, Boehringer Ingelheim, the Canada Foundation for Innovation, Eshelman Institute for Innovation, Genentech, Genome Canada through Ontario Genomics Institute [OGI-196], EU/EFPIA/OICR/McGill/KTH/Diamond, Innovative Medicines Initiative 2 Joint Undertaking [EUbOPEN grant 875510], Janssen, Merck KGaA (aka EMD in Canada and USA), Pfizer, the São Paulo Research Foundation-FAPESP, and Takeda (CIW, CAF, HWO, YL, FMB, JLS, IMG, DHD, ADA). Research reported in this publication was supported in part by the NC Biotechnology Center Institutional Support Grant 2018-IDG-1030 (DHD, ADA), NIH U24DK116204 (DHD, ADA), and NIH 1R44TR001916 (DHD, ADA). This work was supported by the Francis Crick Institute which receives its core funding from Cancer Research UK (CC2037), the UK Medical Research Council (CC2037), and the Wellcome Trust (CC2037); Loulou Foundation Project Grant (11015). Margaux Silvestre is a recipient of the Loulou foundation junior fellowship award (2021). For the purpose of Open Access, the author has applied a CC BY public copyright license to any Author Accepted Manuscript version arising from this submission.

## Acknowledgments

NanoBRET constructs for CDKL5, GSK31Z, and GSK3β were kindly provided to the SGC-UNC team by Promega. TREE*spot* kinase interaction mapping software (http://treespot.discoverx.com) was used to prepare the kinome trees in Figure 3. The authors thank the Diamond Light Source for beamtime (proposal mx28172) as well as the staff of beamline i04 for their guidance.

## Disclosures

Benke – Consultancy for AveXis, Ovid, GW Pharmaceuticals, International Rett Syndrome Foundation, Takeda, Taysha, CureGRIN, GRIN Therapeutics, Alcyone, Neurogene, and Marinus; Clinical Trials with Acadia, Ovid, GW Pharmaceuticals, Marinus and RSRT; all remuneration has been made to his department Axtman – Advisor for Proteic Bioscience Inc.

## Contributions

TAB, SKU and ADA conceived the project. CAF, HWO, YL, FMB, IMG, and DHD designed and synthesized compounds. WR and ANB executed CDKL family thermal shift assays. AC, MS, CIW, JAS, JLS, KD, DHD, NSP, ANB, TAB, SKU, and ADA conducted experiments and analyses. TAB, SKU and ADA drafted the paper, and all contributed to revisions.

## Supplementary Information

### Synthesis of key compounds. Chemistry General Information

Reagents were purchased from reputable commercial vendors and used in accordance with safety data sheets. A rotary evaporator was employed to remove solvent(s) under reduced pressure (*in vacuo*). Thin layer chromatography (TLC) as well as LC–MS were used to monitor reaction progress. The following abbreviations are used in experimental procedures: mmol (millimoles), μmol (micromoles), μL (microliters), mg (milligrams), equiv (equivalent(s)), min (minutes), and h (hours). ^1^H NMR and ^13^C NMR spectra as well as microanalytical data were collected for a key intermediate and final compound **B1** to confirm their identity and evaluate their purity prior to initiating cell- and animal-based studies. ^1^H and ^13^C NMR spectra were collected in DMSO-*d_6_*using Bruker spectrometers. Magnet strength is indicated in each corresponding line listing. Peak positions are included in parts per million (ppm) and calibrated versus the shift of DMSO-*d_6_*; coupling constants (*J* values) are noted in hertz (Hz); and multiplicities are included as follows: singlet (s), doublet (d), pentet of doublets (pd), and multiplet (m).

**Scheme 1.**
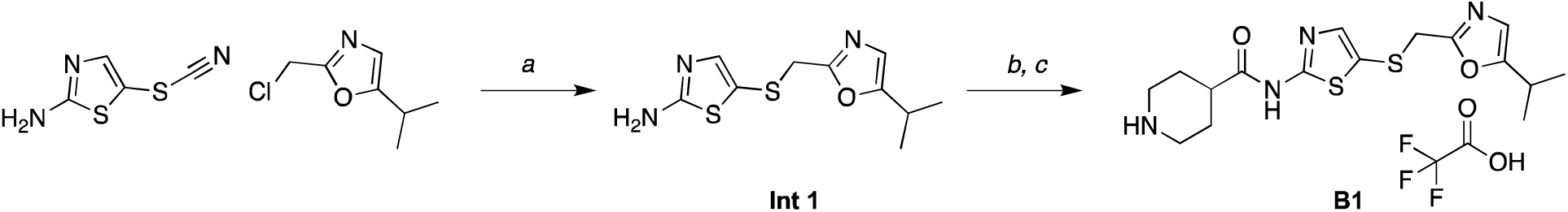
Preparation of compound **B1**. a) EtOH, NaBH_4_, acetone, 80 °C, 1 h, 57% yield; b) HATU, DIPEA, 1-(*tert*-butoxycarbonyl)piperidine-4-carboxylic acid, DMF; c) TFA, CH_2_Cl_2_, 36% yield over two steps.

*5-(((5-isopropyloxazol-2-yl)methyl)thio)thiazol-2-amine (**Int 1**)*. To a flask was added 5- thiocyanatothiazol-2-amine (400 mg, 1.0 equiv, 2.5 mmol) in EtOH (24 mL), followed by addition of NaBH_4_ (183 mg, 1.9 equiv, 4.8 mmol) portion-wise at room temperature. The mixture was stirred for 1 h at room temperature then acetone (12 mL) was added. After 1 hr, a solution of 2- (chloromethyl)-5-isopropyloxazole (412 mg, 1.0 equiv, 2.6 mmol) in EtOH (4mL) was added, and the reaction was heated at 80 °C for 1 h. The resulting mixture was cooled, concentrated *in vacuo*, and then partitioned between EtOAc and brine. The organic phase was separated, dried with MgSO_4_, and concentrated *in vacuo* to give a crude solid. The crude material was triturated with diethyl ether/hexane to provide the desired product 4-(((5-isopropyloxazol-2- yl)methyl)thio)thiazol-2-amine as a red oil (368 mg, 57 %).

*N-(5-(((5-isopropyloxazol-2-yl)methyl)thio)thiazol-2-yl)piperidine-4-carboxamide 2,2,2- trifluoroacetate (**B1**).* To a flask was added 1-(*tert*-butoxycarbonyl)piperidine-4-carboxylic acid (99 mg, 1.1 equiv, 431 μmol) and DIPEA (205 μL, 3 equiv, 1.2 mmol) and HATU (194 mg, 1.3 equiv, 509 μmol) and the flask was stirred at room temperature for approximately 15 min. Next, 5-(((5-isopropyloxazol-2-yl)methyl)thio)thiazol-2-amine (100 mg, 1.0 equiv, 392 μmol) was added and the reaction was stirred at room temperature for 16 h. The reaction mixture was concentrated and purified by column chromatography (SiO_2_, MeOH 0–10% in CH_2_Cl_2_) to yield the desired product as a white solid to yield the intermediate *tert*-butyl 4-((5-(((5- isopropyloxazol-2-yl)methyl)thio)thiazol-2-yl)carbamoyl)piperidine-1-carboxylate. To this intermediate was added 20% TFA and CH_2_Cl_2_ (5 mL) and the reaction stirred for 1 h. The reaction was concentrated *in vacuo* and after addition of MeOH a white precipitate crashed out. The solid was filtered under vacuum to yield the desired product *N*-(5-(((5-isopropyloxazol-2- yl)methyl)thio)thiazol-2-yl)piperidine-4-carboxamide 2,2,2-trifluoroacetate as a white solid (20 mg). The remaining crude material was purified by preparative HPLC (MeOH 10–100% in H_2_O (+0.05% TFA)) to yield the desired product *N*-(5-(((5-isopropyloxazol-2-yl)methyl)thio)thiazol-2- yl)piperidine-4-carboxamide 2,2,2-trifluoroacetate (**B1**) as a white solid (32 mg). Yield over two steps *N*-(5-(((5-isopropyloxazol-2-yl)methyl)thio)thiazol-2-yl)piperidine-4-carboxamide 2,2,2- trifluoroacetate (52 mg, 36 %). ^1^H NMR (400 MHz, DMSO-*d_6_*) δ 7.38 (s, 1H), 6.72 (d, *J* = 1.1 Hz, 1H), 4.03 (s, 2H), 3.21 (d, *J* = 12.8 Hz, 2H), 2.93 – 2.85 (m, 1H), 2.84 – 2.75 (m, 2H), 2.70 (s, 1H), 1.89 (d, *J* = 13.6 Hz, 2H), 1.77 – 1.63 (m, 2H), 1.12 (d, *J* = 6.9 Hz, 6H). ^13^C NMR (214 MHz, DMSO-*d_6_*) δ 172.45, 161.00, 158.76, 158.31, 145.12, 120.91, 119.04, 42.48, 38.65, 34.03, 25.29, 24.83, 20.48. HPLC purity: >95%. HRMS Calculated for [M+H]^+^ C_16_H_23_N_4_O_2_S_2_: 367.1184; observed [M+H]^+^: 367.1248.

**Figure S1.**
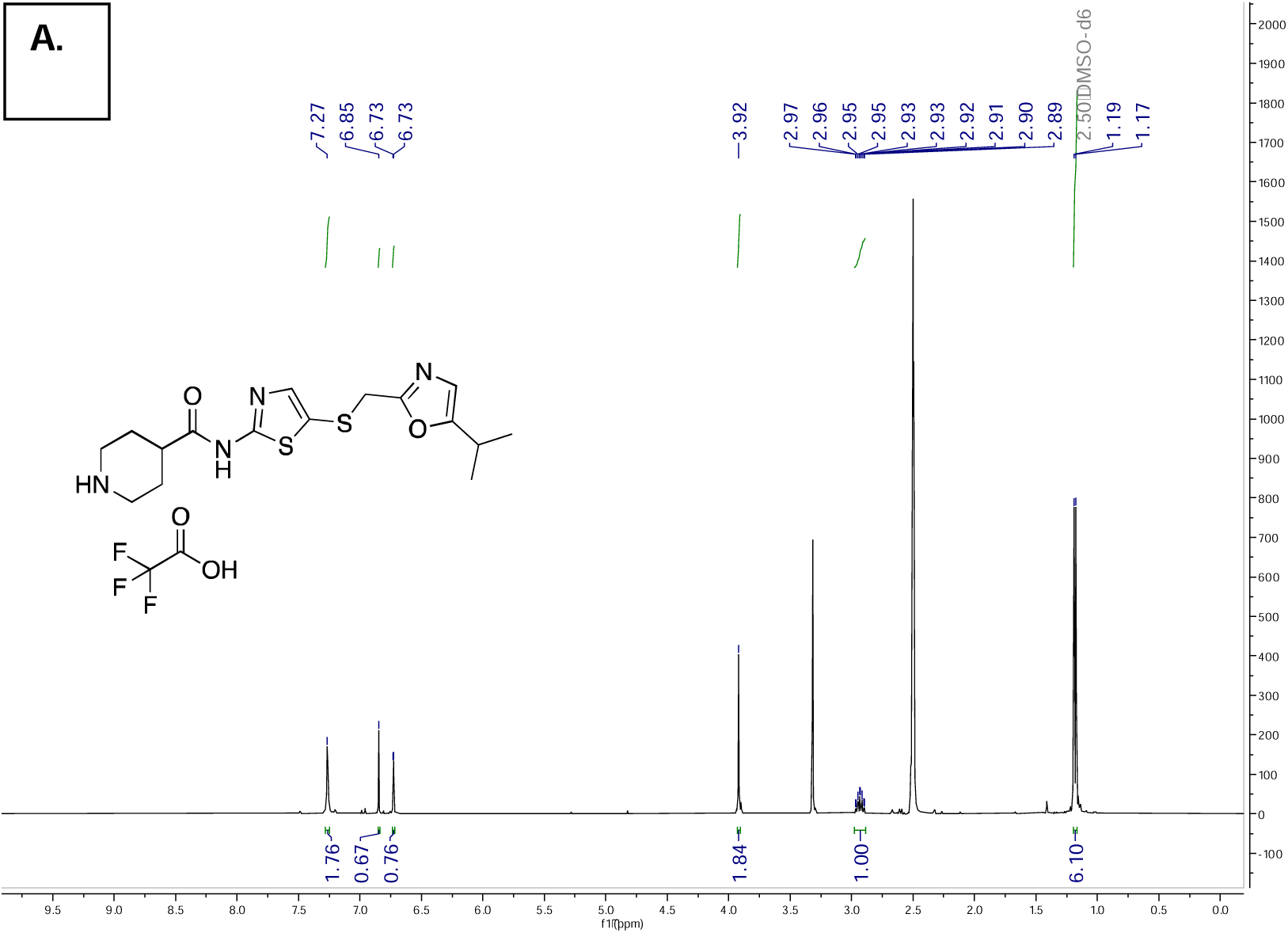
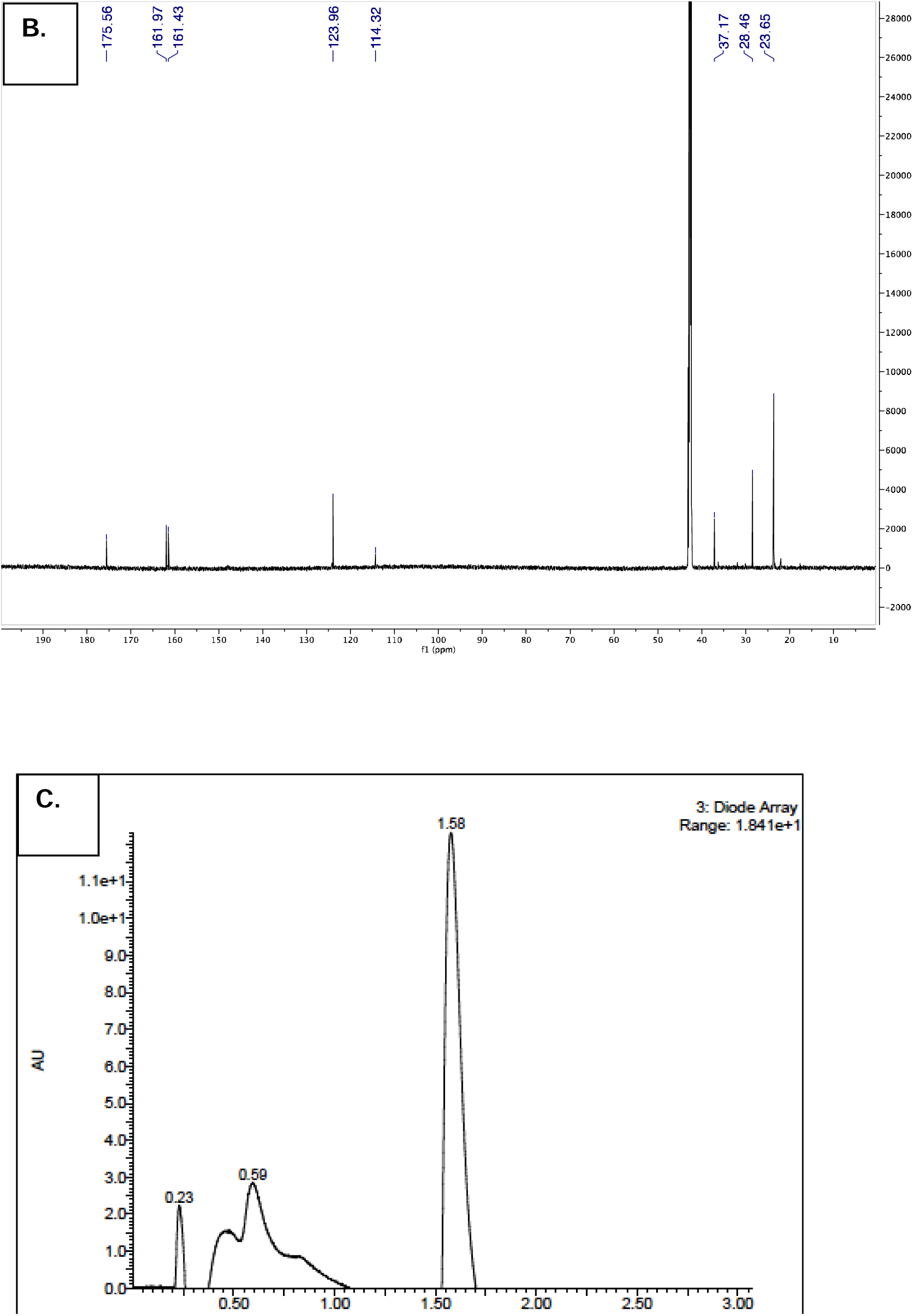
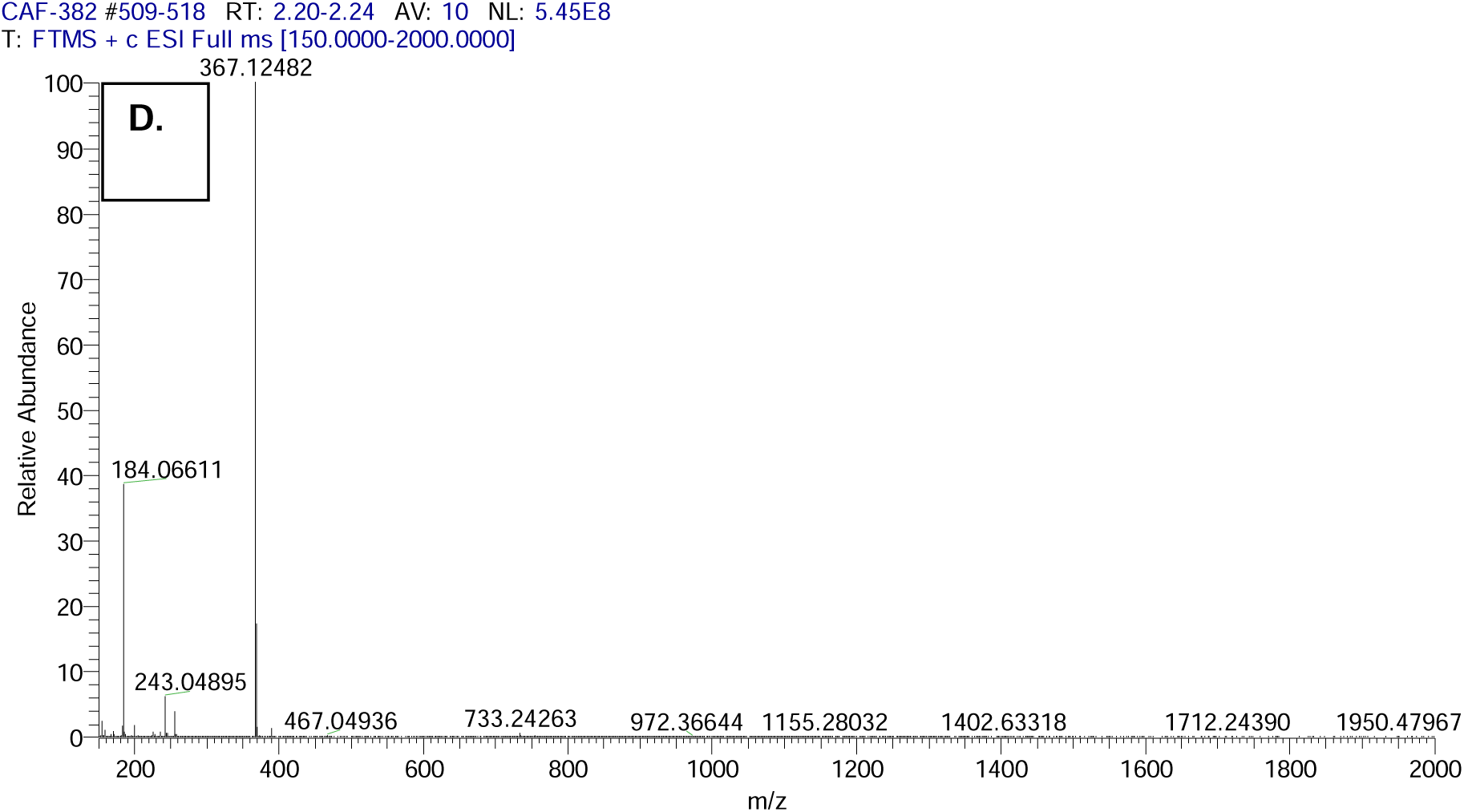
Characterization of N-(5-(((5-isopropyloxazol-2-yl)methyl)thio)thiazol-2-yl)piperidine- 4-carboxamide 2,2,2-trifluoroacetate (**B1**) A) ^1^H Spectra and chemical structure (inset). B) ^13^C Spectra C) HPLC-UV Trace D) HRMS Trace.

**Scheme 2.**
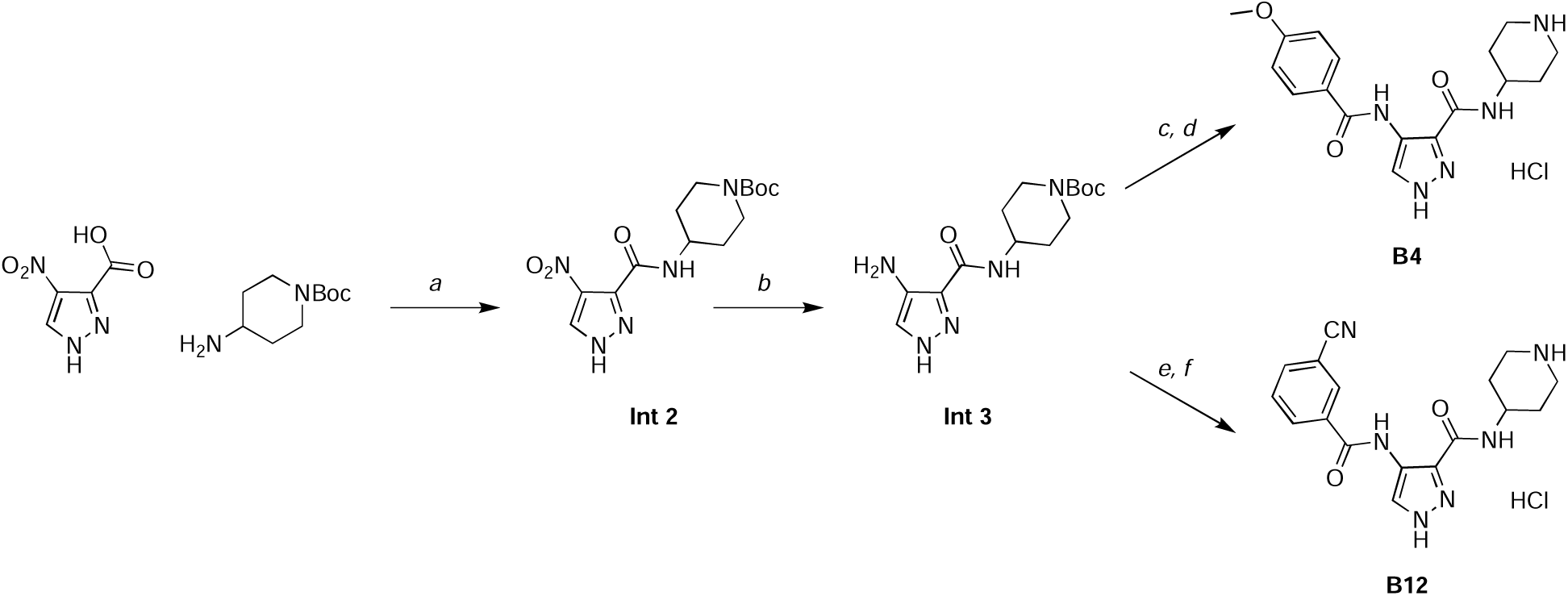
Preparation of compounds **B4** and **B12**. a) Propylphosphonic anhydride (T3P), DIPEA, THF, 0°C, 1 h, 93% yield; b) Palladium on carbon (Pd/C), MeOH, THF, 25°C, 16 h, 99% yield; c) 4-methoxybenzoic acid, HATU, DIPEA, THF, 25°C, 17 h, 72% yield; d) HCl, dioxane, 25°C, 1h, 58% yield; e) 3-cyanobenzoic acid, T3P, DIPEA, THF, 25°C, 16 h; f) HCl, dioxane, 25°C, 1h, 39% yield over two steps.

General procedure for preparation **B4** and **B12**: To a flask was added 4-nitro-1*H*-pyrazole-3- carboxylic acid (1.00 g, 1.0 equiv, 6.4 mmol), 1-Boc-4-aminopiperidine (1.28 g, 1.0 equiv, 6.4 mmol), and DIPEA (4.21 mL, 4.0 equiv, 25.5 mmol) in THF (20 mL). The reaction mixture was cooled to 0°C, and then T3P (6.08 mL, 1.5 equiv, 9.5 mmol) was slowly added. After stirring for 1 h at 0°C, the reaction mixture was concentrated *in vacuo*, and then partitioned between EtOAc and brine. The organic phase was separated, dried with MgSO_4_, and concentrated *in vacuo* to give a crude solid. The crude material was purified using an automated purification system (SiO_2_) 90%/10% hexanes/EtOAc to 100% EtOAc to give *tert*-butyl 4-(4-nitro-1*H*-pyrazole-3- carboxamido)piperidine-1-carboxylate (**Int 2**) as an amorphous solid (2.00 g, 93% yield).

Palladium on carbon (5%, 31.3 mg, 0.1 equiv, 0.295 mmol) was added to *tert*-butyl 4-(4- nitro-1*H*-pyrazole-3-carboxamido)piperidine-1-carboxylate (**Int 2**, 1.0 g, 1.0 equiv, 2.95 mmol) in a mixture of methanol (10 mL) and THF (3.0 mL). The solution was placed under an atmosphere of H_2_ at room temperature. After 16h, the solution was filtered through SiO_2_ and the eluent was concentrated to afford *tert*-butyl 4-(4-amino-1*H*-pyrazole-3-carboxamido)piperidine- 1-carboxylate (**Int 3**) as an amorphous solid (900 mg, 99% yield).

To a flask was added *tert*-butyl 4-(4-amino-1*H*-pyrazole-3-carboxamido)piperidine-1- carboxylate (**Int 3**, 1 equiv) and the corresponding carboxylic acid (1.2 equiv) and DIPEA (4.0 equiv) in THF (0.5 M) at room temperature. The reaction mixture was then treated slowly with T3P (3.0 equiv). After stirring for 16–17 h, the reaction mixture was concentrated *in vacuo*, and then partitioned between EtOAc and brine. The organic phase was separated, dried with MgSO_4_, and concentrated *in vacuo* to give a crude solid. The crude material was purified using an automated purification system (SiO_2_) 90%/100% hexanes/EtOAc to 100% EtOAc) to give an intermediate, which was stirred in HCl/dioxane (3.3 mL) for 1 h at room temperature. Solvent was removed and the crude material was purified using an automated purification system (SiO_2_) 90%/100% hexanes/EtOAc to 100% EtOAc) to give the desired product (**B4** or **B12**) as an amorphous solid (39–42% yield over 2 steps).

*4-(4-methoxybenzamido)-N-(piperidin-4-yl)-1H-pyrazole-3-carboxamide* (**B4**) ^1^H NMR (850 MHz, Methanol-*d*_4_) δ 8.30 (s, 1H), 7.89 (d, *J* = 8.8 Hz, 2H), 7.07 (d, *J* = 8.8 Hz, 2H), 4.22 (tt, *J* = 10.8, 4.1 Hz, 1H), 3.89 (s, 3H), 3.48 (dt, *J* = 13.4, 3.9 Hz, 2H), 3.18 (td, *J* = 12.7, 3.1 Hz, 2H), 2.22 (dd, *J* = 14.3, 3.9 Hz, 2H), 1.96 – 1.85 (m, 2H). ^13^C NMR (214 MHz, Methanol-*d*_4_) δ 165.66, 165.58, 164.53, 134.00, 130.01, 126.71, 124.57, 121.57, 115.24, 56.06, 45.26, 44.25, 29.54. ^1^H NMR purity: >95%. HRMS Calculated for [M + H]^+^ C_17_H_22_N_5_O_3_: 344.1723; observed [M+H]^+^: 344.1717.

**Figure S2.**
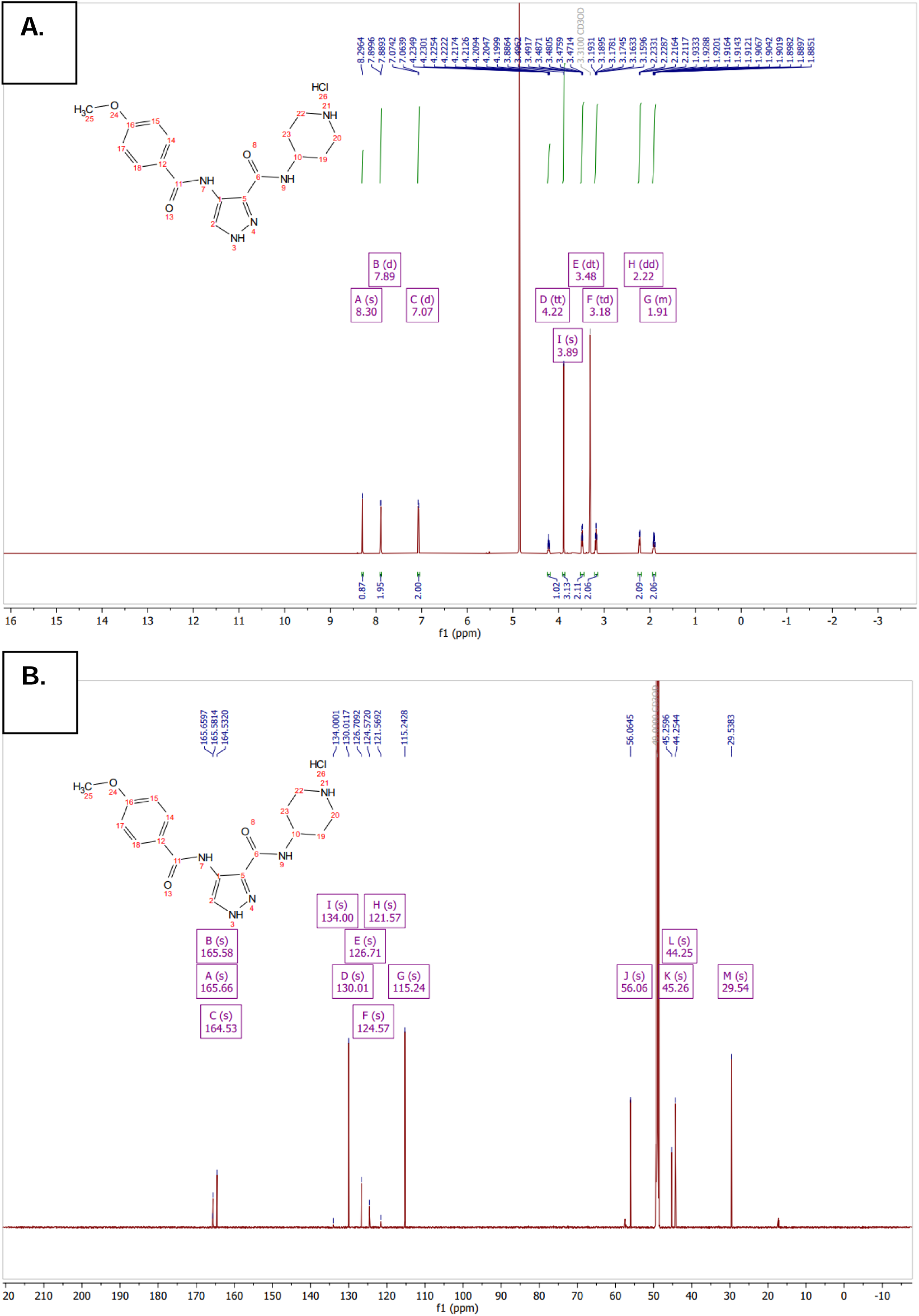
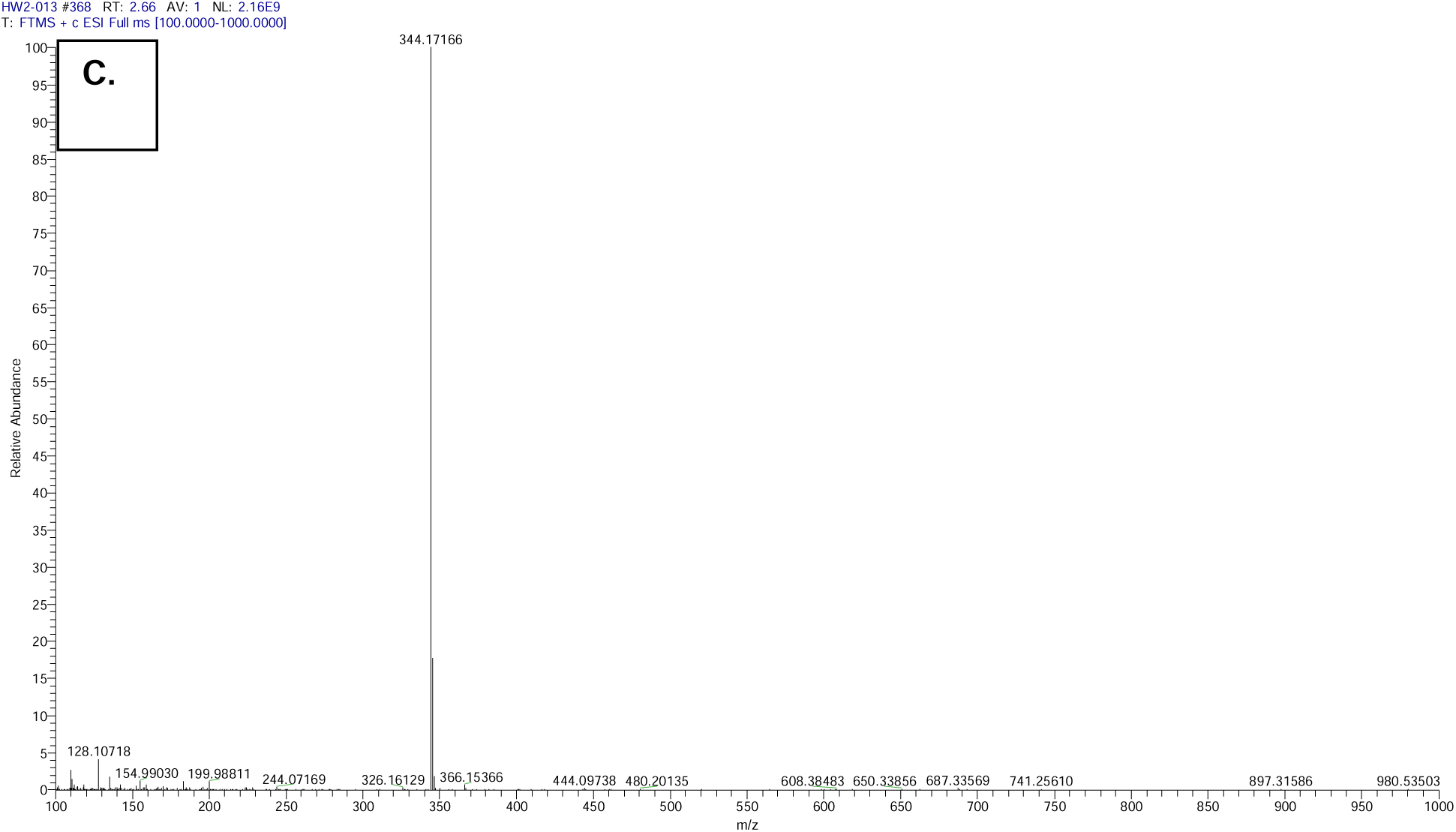
Characterization of 4-(4-methoxybenzamido)-N-(piperidin-4-yl)-1H-pyrazole-3-carboxamide (**B4**) A) ^1^H Spectra and chemical structure (inset). B) ^13^C Spectra C) HRMS Trace.

*4-(3-cyanobenzamido)-N-(piperidin-4-yl)-1H-pyrazole-3-carboxamide*(**B12**) ^1^H NMR (850 MHz, Methanol-*d*_4_) δ 8.33 (s, 1H), 8.26 (t, *J* = 1.7 Hz, 1H), 8.22 (dt, *J* = 7.9, 1.5 Hz, 1H), 7.97 (dt, *J* = 7.7, 1.4 Hz, 1H), 7.75 (t, *J* = 7.8 Hz, 1H), 4.22 (tt, *J* = 10.9, 4.1 Hz, 1H), 3.49 (dt, *J* = 14.1, 3.5 Hz, 2H), 3.18 (td, *J* = 12.9, 3.1 Hz, 2H), 2.22 (dd, *J* = 14.2, 3.7 Hz, 2H), 1.95 – 1.88 (m, 2H). ^13^C NMR (214 MHz, Methanol-*d*_4_) δ 165.57, 163.51, 136.50, 136.14, 134.35, 132.50, 131.84, 131.32, 124.01, 122.04, 118.88, 114.39, 45.33, 44.29, 29.53. ^1^H NMR purity: >95%. HRMS Calculated for [M + H]^+^ C_17_H_19_N_6_O_2_: 339.1569; observed [M+H]^+^: 339.1563.

**Figure S3.**
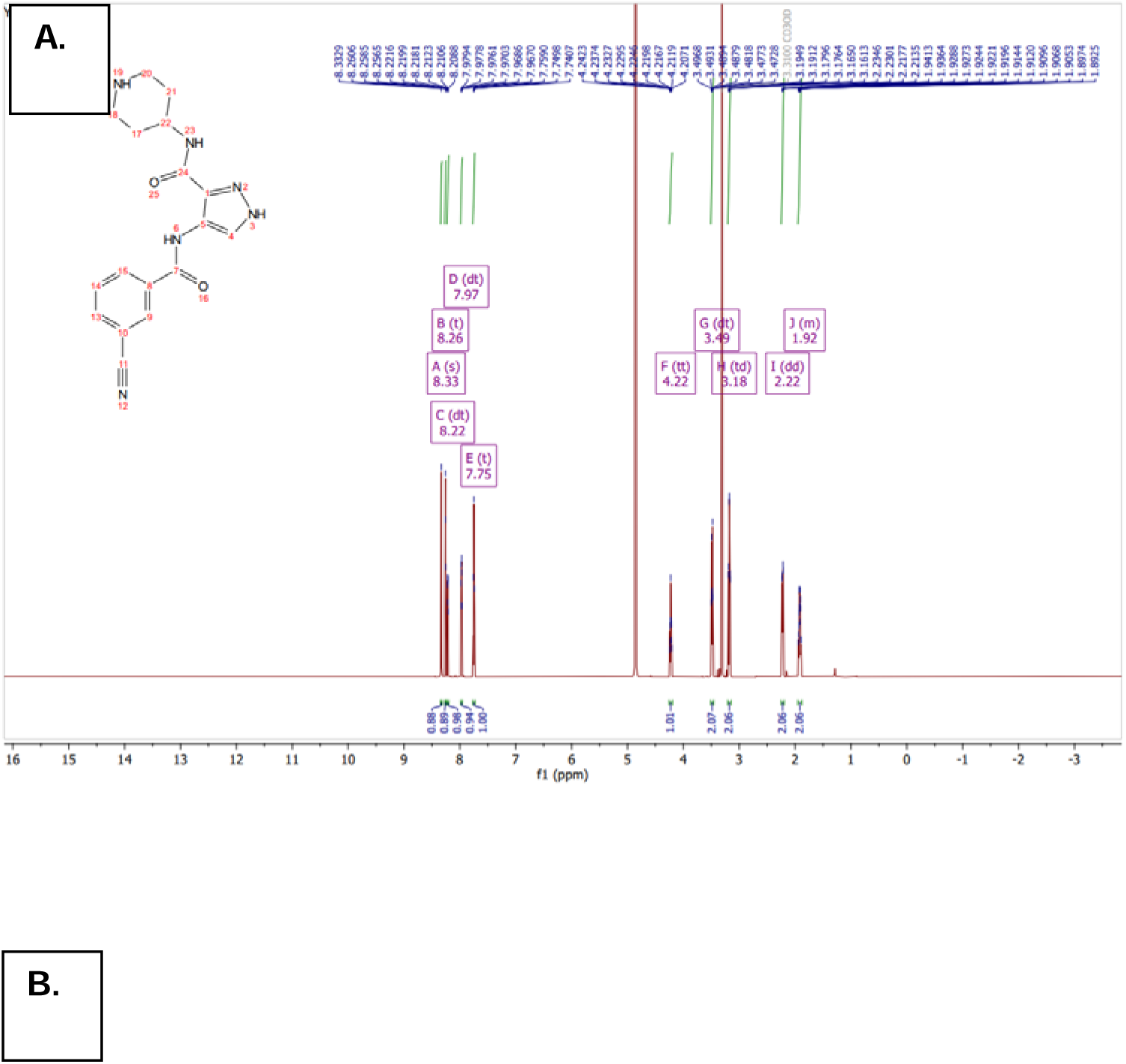
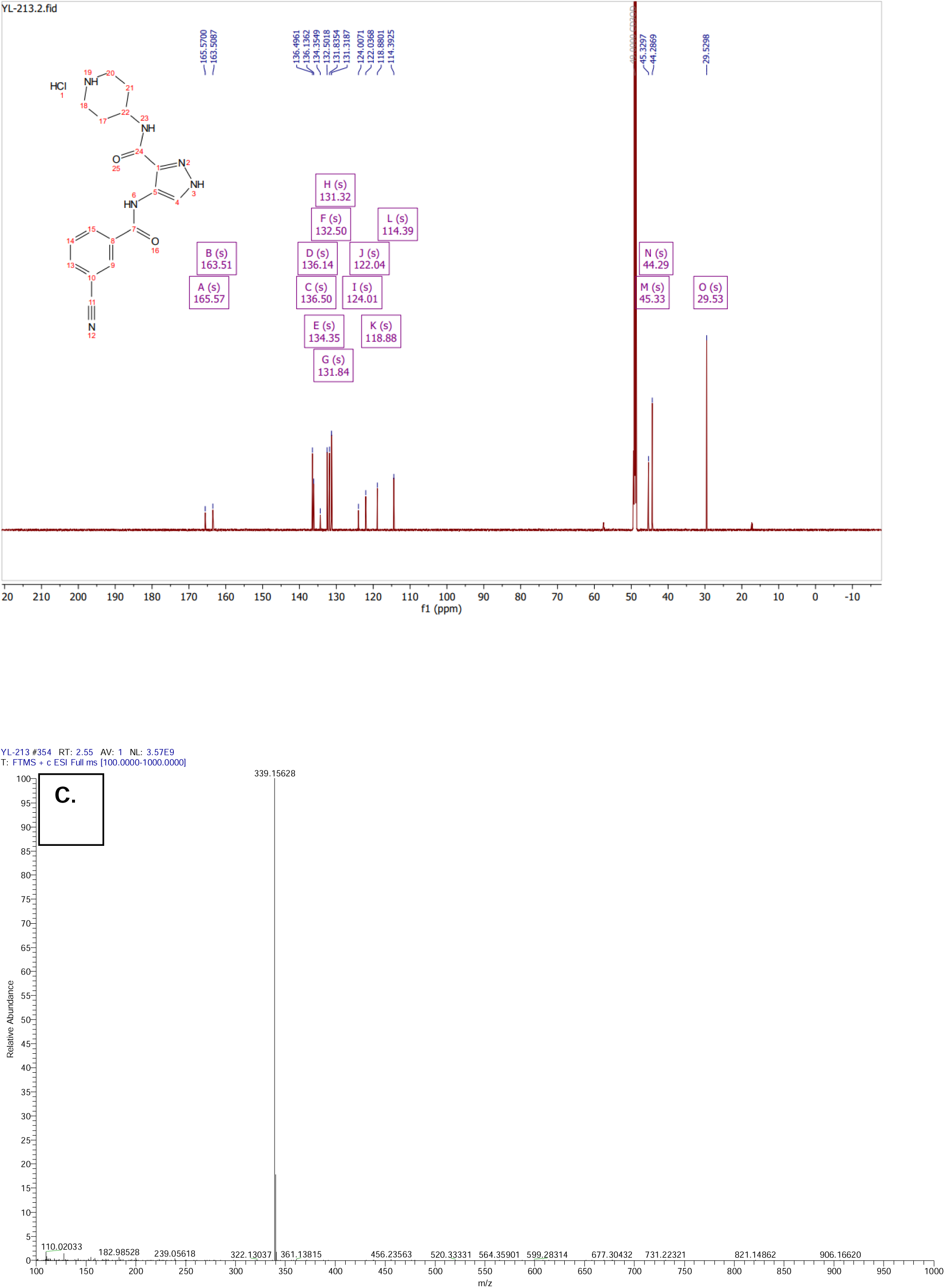
Characterization of 4-(3-cyanobenzamido)-N-(piperidin-4-yl)-1H-pyrazole-3-carboxamide (**B12**) A) ^1^H Spectra and chemical structure (inset). B) ^13^C Spectra C) HRMS Trace.

**Table S1.** NanoBRET data corresponding to compounds selected for initial study.

| **Compound** | **Parent** | **CDKL5 NB IC<sub>50</sub> (nM)** | **GSK3β NB IC<sub>50</sub> (nM)** | **fold** |
| --- | --- | --- | --- | --- |
| A01 | AT-7519 | 363 | >10,000 | 27.5 |
| A02 | AT-7519 | 178 | 4135 | 23.2 |
| A03 | AT-7519 | 19 | 404.8 | 21.3 |
| A04 | SNS-032 | 108 | 5688 | 52.7 |
| A05 | AT-7519 | 12 | 308.2 | 25.7 |
| A06 | SNS-032 | 221 | 7787 | 35.2 |
| A07 | AT-7519 | 6 | 192 | 32.0 |
| A08 | AT-7519 | 56 | 1374 | 24.5 |
| A09 | AT-7519 | 21 | 481 | 22.9 |
| A10 | SNS-032 | 424 | >10,000 | 23.6 |
| A11 | AT-7519 | 9 | 3699 | 411.0 |
| A12 | AT-7519 | 20 | 524.8 | 26.2 |
| B01 | SNS-032 | 8 | >10,000 | 1250.0 |
| B02 | SNS-032 | 190 | 9100 | 47.9 |
| B03 | SNS-032 | 10 | >10,000 | 1000.0 |
| B04 | AT-7519 | 14 | 553.9 | 39.6 |
| B05 | SNS-032 | 388 | 8000 | 20.6 |
| B06 | AT-7519 | 155 | 8193 | 52.9 |
| B07 | AT-7519 | 36 | 3256 | 90.4 |
| B08 | AT-7519 | 161 | >10,000 | 62.1 |
| B09 | AT-7519 | 37 | 3627 | 98.0 |
| B10 | AT-7519 | 90 | 4945 | 54.9 |
| B11 | AT-7519 | 141 | 3101 | 22.0 |
| B12 | AT-7519 | 69 | 3631 | 52.6 |
| C01 | SNS-032 | 323 | >10,000 | 31.0 |

**Figure S4.**
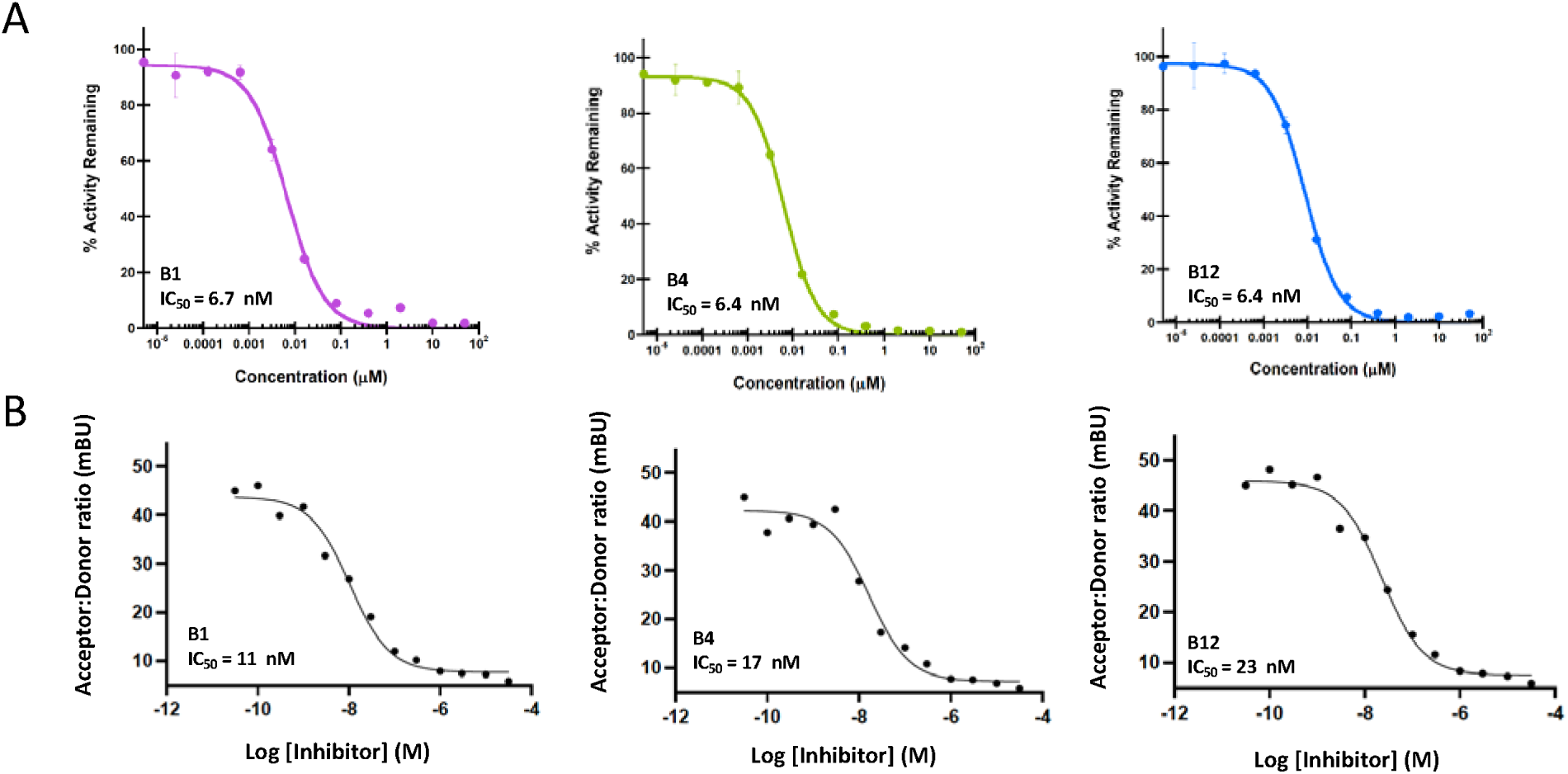
Curves corresponding with CDKL5 affinity measurements in Figure 2. (A) Curves generated in CDKL5 split-luciferase assay for B1, B4, and B12. (B) Curves generated in CDKL5 NanoBRET assay for B1, B4, and B12.

**Figure S5.**
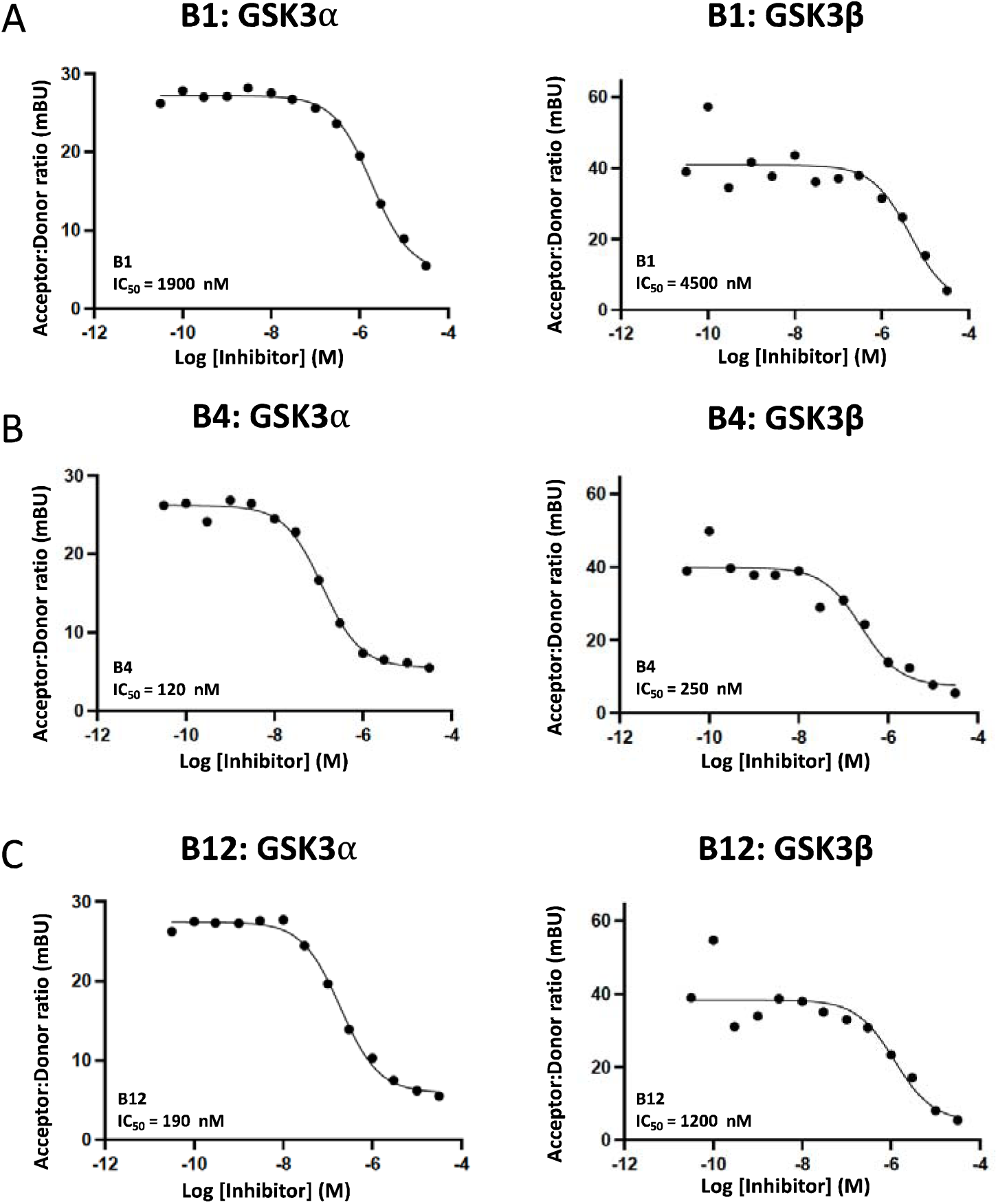
Curves corresponding with GSK3 NanoBRET measurements in Figure 2. (A) Curves generated in GSK31Z and GSK3β NanoBRET assays for B1. (B) Curves generated in GSK31Z and GSK3β NanoBRET assays for B4. (C) Curves generated in GSK31Z and GSK3β NanoBRET assays for B12.

**Figure S6.**
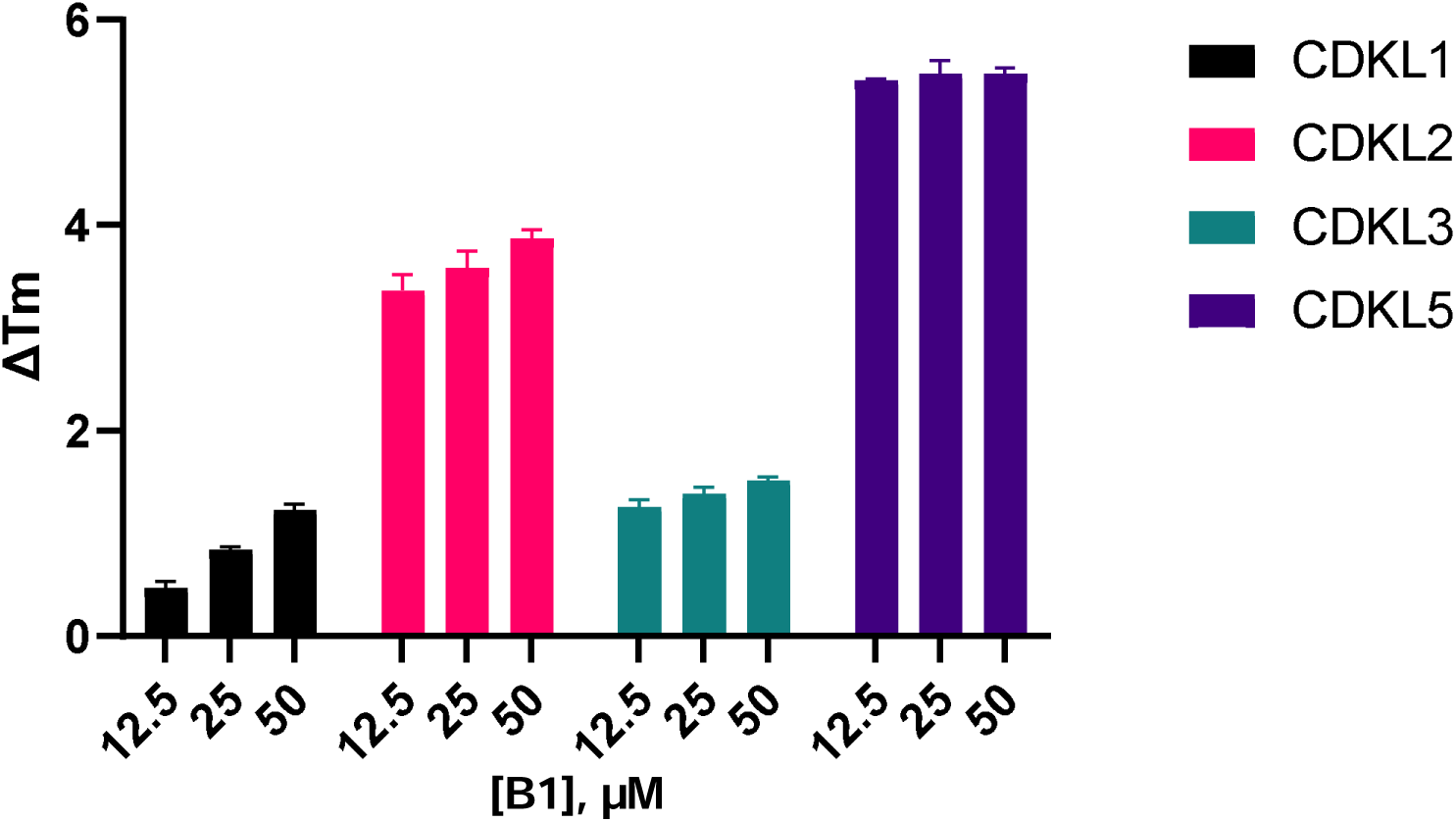
CDKL family selectivity evaluation via thermal shift assays.

**Figure S7.**
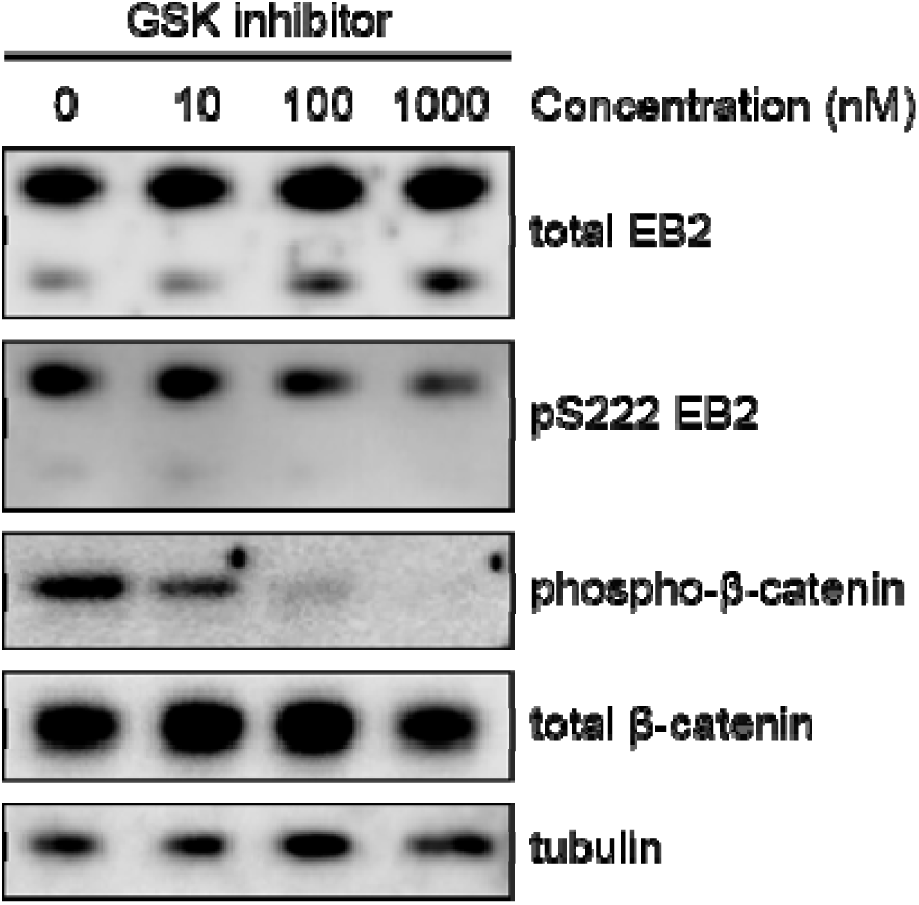
Phosphorylation of β-catenin, a substrate of GSK, is reduced upon treatment with a GSK inhibitor CHIR 99021 (Tocris) but EB2 phosphorylation is not changed. Western blot showing expression of total EB2, pS222 EB2, total β-catenin, phospho-β-catenin and tubulin in DIV14-15 rat primary neurones after an hour treatment with a GSK inhibitor.

**Figure S8.**
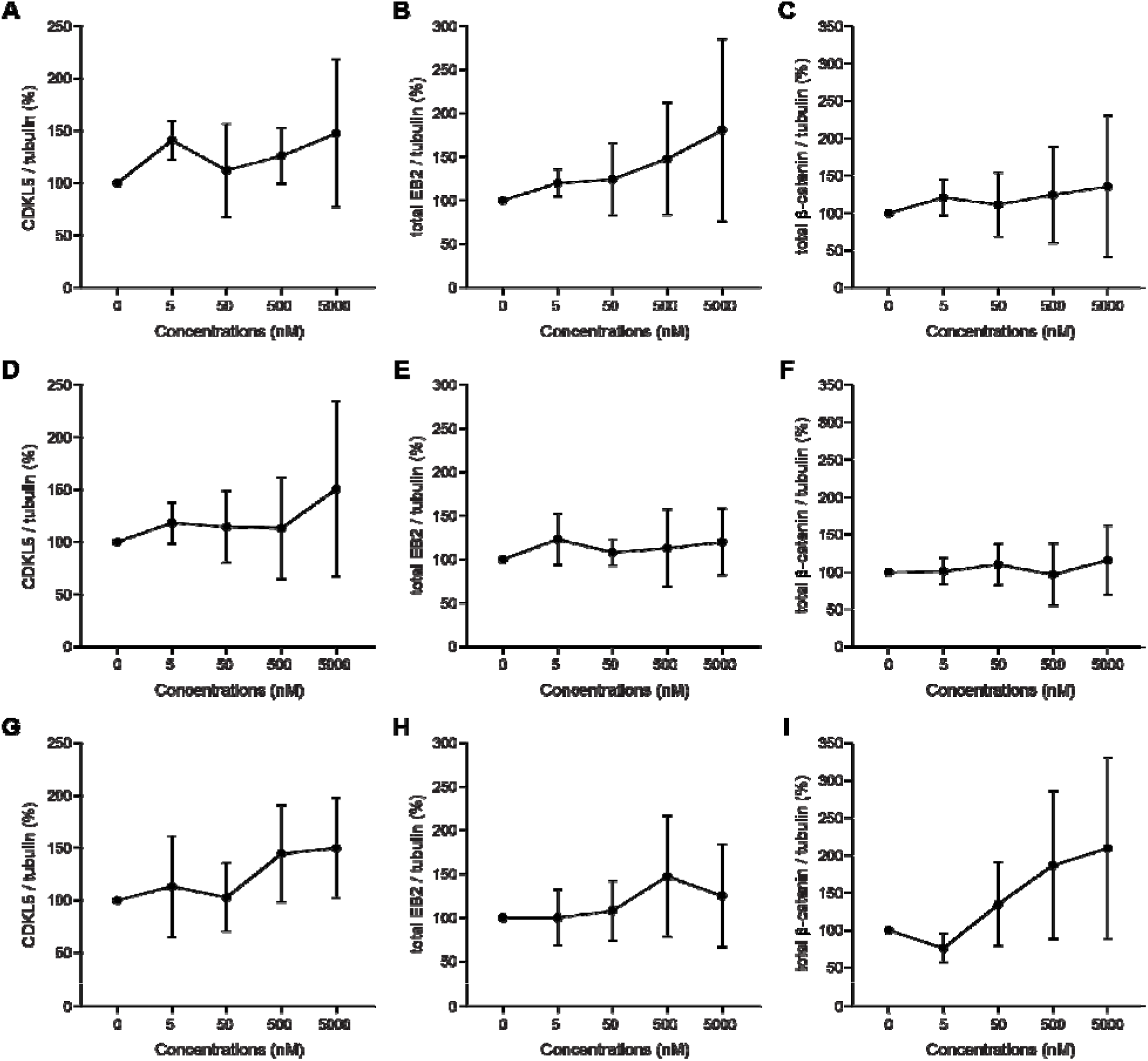
CDKL5, total EB2 and β-catenin expression in rat primary neurons after treatment with CAF-382, HW2-013 and LY-213 compounds. A, D, G, Quantification of CDKL5 expression in DIV14-15 rat primary neurones after an hour treatment with different concentrations of CAF-382, HW2-013 and LY-213 compounds respectively. Each concentration was compared to the control using a one-way ANOVA test. N = 3 biological replicates with 2 repetitions. Error bars are SD. B, E, H, Quantification of total EB2 expression in DIV14-15 rat primary neurones after an hour treatment with different concentrations of CAF-382, HW2-013 and LY-213 compounds respectively. Each concentration was compared to the control using a one-way ANOVA test. N = 3 biological replicates with 2 repetitions. Error bars are SD. C, F, I, Quantification of total β-catenin expression in DIV14-15 rat primary neurones after an hour treatment with different concentrations of CAF-382, HW2-013 and LY-213 compounds respectively. Each concentration was compared to the control using a one-way ANOVA test. N = 2 biological replicates with 2 repetitions. Error bars are SD.

**Figure S9.**
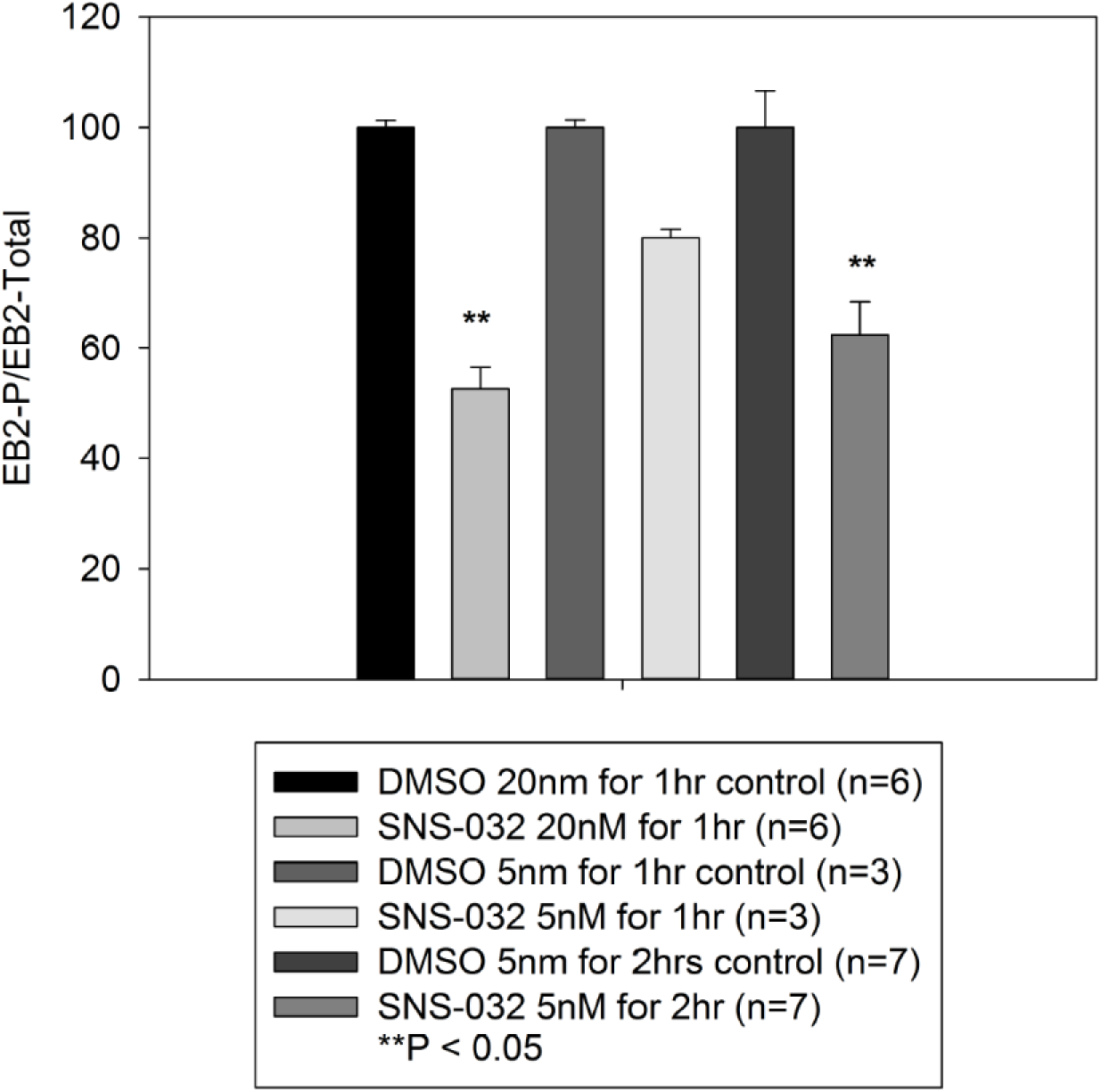
SNS-032 (Selleck Chemicals) inhibits phosphorylation of EB2 in rat hippocampal slices in a time and concentration-dependent fashion. Effect of SNS-032 is compared to control slices with similar concentration of DMSO vehicle under three different conditions. Compare to Figure 6.

**Figure S10.**
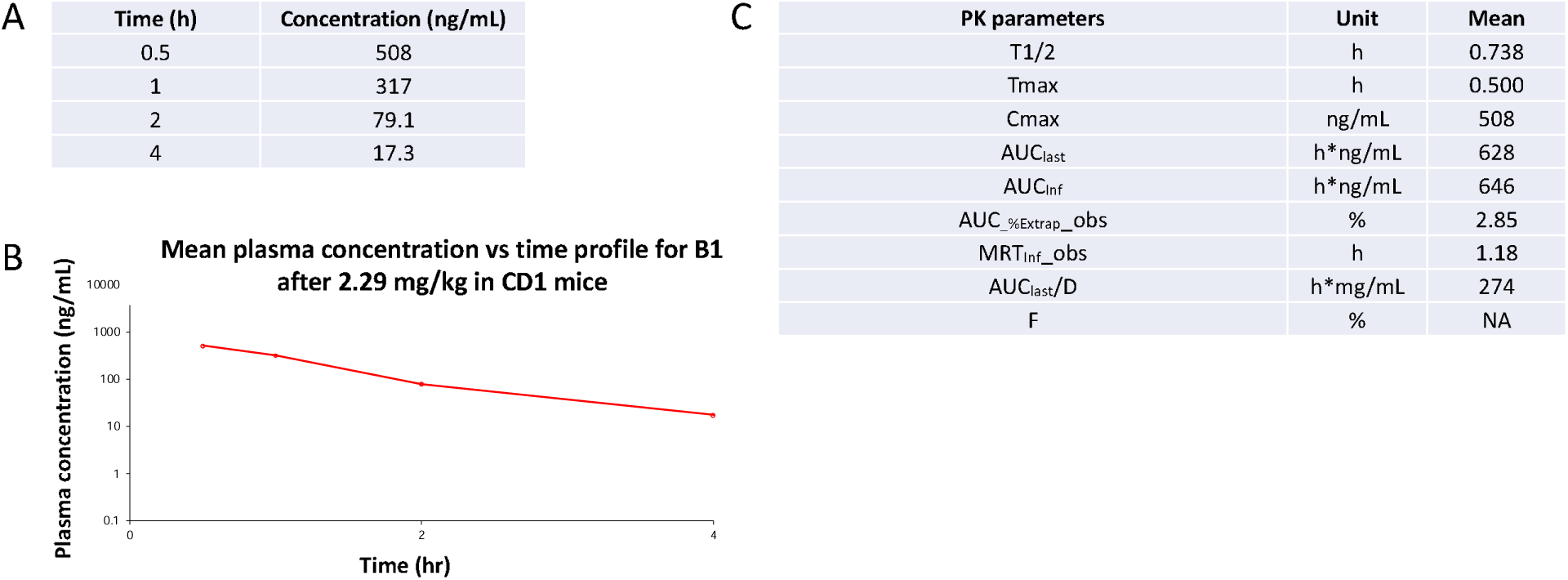
Snapshot pharmacokinetic (PK) results for B1. (A) Plasma concentration measurements at timepoints post-administration. (B) Plot of mean plasma concentration over the time course of the PK study. (C) Summary of B1 PK properties (n = 2).

**Table S2.** Brain exposure results for B1. (A) Brain/plasma concentration-time data after 2.29 mg/kg dose. (B) Brain/plasma concentration-time data after 7.63 mg/kg dose. *One animal artificially drove up blood/plasma ratio. N = 3 per dose.

| A | IP dose: 2.29 mg/kg |  |  |  | B | IP dose: 7.63 mg/kg |  |  |  |
| --- | --- | --- | --- | --- | --- | --- | --- | --- | --- |
|  | Time (h) | Brain Conc (ng/g) | Plasma (ng/mL) | Ratio (Brain/plasma) |  | Time (h) | Brain Conc (ng/g) | Plasma (ng/mL) | Ratio (Brain/plasma) |
|  | 1 | 15.6 | 340 | 0.0459 |  | 1 | 158 | 750 | 0.210* |

## Notes

### Competing Interest Statement

The authors have declared no competing interest.

### Summary of Updates

Updates to Figure 4. Summary of initial reviews.

