## Supplementary material for "Discovery and characterization of a specific inhibitor of serine-threonine kinase cyclin dependent kinase-like 5 (CDKL5) demonstrates role in hippocampal CA1 physiology": reviewer comments

### Benke, Tim

**From:**
**Sent:** Thursday, June 8, 2023 9:48 AM
**To:** Benke, Tim
**Subject:** Reviews for your submission to eLife: 29-03-2023-RA-eLife-88206

[External Email - Use Caution]

Dear Dr Benke,

Thank you for submitting your article "Discovery and characterization of a specific inhibitor of serine-threonine kinase cyclin dependent kinase-like 5 (CDKL5) demonstrates role in hippocampal CA1 physiology" for consideration by eLife. Your article has been reviewed by 3 peer reviewers, and the evaluation has been overseen by a Reviewing Editor and Amy Andreotti as the Senior Editor.

The reviewers have discussed their reviews with one another, and the Reviewing Editor has drafted this to help you prepare a revised submission.

----------

Essential Revisions (for the authors):

1) Please further describe the screening procedure that led to the reported selective inhibitors (e.g., size of the screened libraries, description of the binding and orthogonal assays, reasons to use myelin basic protein as a substrate for CDKL5 in the in vitro kinase assays, etc.).

2) Please discuss potential limitations to using the reported inhibitors based on the seemingly limited brain penetration.

3) Please revise figures, to clearly show statistically significant differences when detected, add legends, improve description in Figures' captions, etc.

4) On the basis of the reviewers' comments. Please further discuss the statement "At issue, is whether observations of CDKL5 activity, either through genetic manipulations or interference with the kinase, are specific and not potentially subsumed by the activity of other kinases. Given the reduction of phosphorylation of the specific substrate, EB2, this is less likely here".

----------

eLife's editorial process also produces an assessment by peers designed to be posted alongside a preprint for the benefit of readers.

eLife assessment

This important study reports selective CDKL5 inhibitors that may be of high interest to investigate the role of this kinase in disease (particularly, in CDKL5 deficiency disorder) and to address unsolved issues of inconsistency in the phenotypic characterization of CDKL5-deficient knockout mice. The evidence supporting the conclusions is convincing, with rigorous biochemical, in vitro and ex vivo assays. The work will be of interest to cell and medical biologists and epileptologists working in the fields of epilepsy and neural excitation.

Reviewer #1 (Public Review):

Castano et al. report a screening to search for selective CDKL5 inhibitors. After profiling an extensive library of selective cyclin-dependen kinase inhibitors, the authors synthesized and characterized high-affinity selective inhibitors of CDKL5. Kinome-wide studies were performed to verify selectivity. Preliminary PK studies were realized in rodents, including the determination of total brain/plasma ratio associated with two dose levels and microsomal stability. When applied directly to rat hippocampal brain slices, one of the inhibitors (CAF-382) reduced post-synaptic function of AMPA-type glutamate receptors dose-dependently, and also reduced hippocampal long-term potentiation. CAF-382 could be a valuable tool to further investigate the role of CDKL5 in disease, although some potential applications may be limited by the seemingly low brain bioavailability of this compound.

The conclusions of this paper are in general terms well supported by data, but some aspects of the discussion of the results could be extended.

Reviewer #1 (Recommendations for the authors):

In the introduction, when stating: "Epilepsy in CDD does not respond to any antiseizure medications[11]; a newly approved therapy provides modest improvements[12]", the identity of this newly approved therapy should be revealed.

Please indicate the source(s) of reactants used for synthesis.

Even if some of the reported data were obtained through an outsource service (e.g., the studies on kinome-wide selectivity) the details of the assay should be exhaustively described under the methods section to ensure reproducibility. Although a reference (ref. 43) has been included related to kinase inhibition selectivity, the authors are not from the same organization that has measured selectivity here, so I believe the details on the methodology should be made explicit.

Regarding the preliminary PK study, I am curious about how tmax was estimated at 0.5 hours considering this is the earliest sampling point. Also, abbreviations used in Fig. S10 should be defined and data should be presented with dispersion. How was the administered dose chosen? I am not sure about reporting Cmax and tmax with such a limited number of data points and no earlier sampling times.

How were the doses used for the brain/plasma partitioning chosen?

Details on the housing and feeding of the laboratory animals at The Francis Crick Institute should be provided.

What is the total number of compounds included in the extensive libraries of SNS-032 and AT-7519 analogs?

Whereas the S10 score is defined in the Figure 3 caption, it would be a good idea to include its definition in the main text, possibly in the Method section.

Have the authors obtained data on the unbound fraction of B1?

I am not convinced by the argument that, since B1 reduces the phosphorylation of the specific substrate EB2, interference with other kinases expressed in the brain might not be at play here. Please elaborate further.

The authors indicate that brain penetration of B1 is rather low. Do they see this as an obstacle to using their inhibitor to further study the role of CDKL5 in CDD using in vivo assays? What about determining the kpuu, which is much more relevant pharmacologically speaking? (the shift in potency from biochemical to cell-based assays also suggests a permeability problem). Haven´t the authors considered a hit-to-lead campaign to improve B1 PK profile?

The potential utility of B1 could be further highlighted towards the concluding section of the manuscript.

I suggest indicating the statistically significant differences in the means in Figure 6 graphically. Can the authors explain the apparent lack of a clear dose-response behavior?

Seemingly, the authors have not observed their code to indicate different levels of statistical significance in Figure 7.

What conclusion(s) can the authors draw from the similar microsomal stability observed with and without NADPH? Are the differences between 0 and 30 min statistically significant? Please provide a wider interpretation of the results.

Reviewer #2 (Public Review):

In the present study, Castano et al. discovered a chemical inhibitor that is specifically effective against the kinase activity of CDKL5 and applied it in the in vitro and the brain slice culture to reveal the acute effects of the loss of function (LOF) of CDKL5. LOF has been modeled in gene knockout mice, but these are loss-of-function models with the added developmental time effects of the absence of CDKL5 from developmental stages. The present authors' approach is the fastest timescale study to date, examining CDKL5 LOF effects in seconds to minutes.

The authors showed that chemical inhibition of CDKL5 kinase activity suppresses postsynaptically derived LTP in rat brain slice experiments, indicating that the previously controversial results of CDKL5 LOF on LTP in knockout mice and rats are possibly due to combined effects of the loss of the kinase and compensation by other factors.

The authors employed state-of-the-art methodologies and presented their data clearly and convincingly.

Reviewer #2 (Recommendations for the authors):

I have several questions to clarify some descriptions.

1. Selectivity score
Figure 3 and the corresponding description of the selectivity score - the meaning of these is hard to understand. Do the dots in kinome tree diagrams represent 403 kinases? Which dot is CDKL5? Why were red circles drawn only on the CMGC family? The authors described that the S10 score is a way to express selectivity that corresponds with the percent of the kinases screened that bind with a PoC value <10 and indicated 0.017, 0.02, and 0.01 for B1, B4, and B12 respectively. But it is hard to interpret the significance of these values. Can you provide some scale or perspective and let us know how selective these values are in terms of the kinome? Why do some kinases have two IC50 (nM) values?

2. Figure 7B
Please add legends for 3 traces.

3. Lines 653-656
"At issue, is whether observations of CDKL5 activity, either through genetic manipulations or interference with the kinase, are specific and not potentially subsumed by the activity of other kinases. Given the reduction of phosphorylation of the specific substrate, EB2, this is less likely here."
It is hard to understand the intent of these sentences.

Reviewer #3 (Public Review):

In this manuscript, Castano et al generate and test a small molecule inhibitor of CDKL5, an X-linked kinase whose loss-of-function is the cause of a severe neurodevelopmental disorder. Since the current knowledge of CDKL5 functions mainly rely on genetic models it is still unclear which effects are caused directly by CDKL5 loss and which can be ascribed to indirect effects. A specific inhibitor would therefore be an important tool for the field.

Castano and colleagues therefore tested a panel of twenty kinase inhibitors for their capacity to block phosphorylation of a EB2, a bona fide CDKL5 substrate, in rat neurons. Among the three that could inhibit EB2 phosphorylation at low concentrations, one was found to inhibit CDKL5 while not affecting GSK3 kinases, which share significant homology to CDKL5. Considering that genetic studies have previously linked CDKL5 to excitatory synaptic transmission, acute hippocampal slices were exploited to test the consequences of CDKL5 inhibition. While CDKL5 loss in the past was found to affect both AMPA- and NMDA-Rs, the small molecule-based inhibition affected only AMPA-R responses at the post-synaptic level. Since pharmacokinetic analyses showed that the inhibitor has a low capacity for brain penetration the molecule remains limited for testing the acute inhibition of CDKL5 in vitro and ex vivo. Such a tool represents an important aspect in the CDKL5 field and the findings suggesting a direct role of CDKL5 in regulating AMPA-R functions are interesting. However, the manuscript could be improved to render it more readable.

The description of the binding and orthogonal assays, which are the basis for the selection of the small molecule inhibitor, is not straightforward to understand for non-expert readers and could be improved.

While the in vitro and ex vivo assays are well presented, it is not clear why the myelin basic protein is used as a substrate for CDKL5 in the in vitro kinase assays. Does this protein contain a CDKL5 consensus site?

Reviewer #3 (Recommendations for the authors):

The first part of the paper describing binding and orthogonal assays should be improved to render the read easier. For non-expert readers, it is impossible to really understand the specificity of the inhibitor, which is a central part of the manuscript. It is also not clear why the authors start with a panel of 25 inhibitors and test 20 in the rat neurons.

While the work in general is well presented it could be improved by being updated and, in a few cases, more precise.

Line 170: "epilepsy in CDD does not respond to any seizure medication" - that is a bit overstated.

Line 182: I suggest citing Terzic and colleagues' work from 2021 showing that CDKL5 is required also during adulthood.

Line 192-197: I suggest including De Rosa et al's work from 2023 suggesting a role of CDKL5 in GABAA-R mediated inhibition through the interaction with gephyrin.

Line 590: the work by Kontaxi et al has been published.

Kinase assay: it is not clear whether full-length CDKL5, and which isoform, was used for the in vitro kinase assay. Furthermore, it is not clear why myelin basic protein was used as a substrate - does it contain the CDKL5 consensus site (which is by the way never mentioned in the paper)?

----------

POSTING PUBLIC REVIEWS: If you plan to resubmit to eLife, please confirm when you are ready for us to proceed with us posting the eLife assessment and Public Reviews to accompany your preprint (and please include author responses if you would like them to be posted at the same time).

If there are clear factual errors in these sections that preclude public posting, please let us know. Any sections marked as "Essential Revisions" or "Recommendations for the authors" will NOT be posted to the preprint.

----------

Information about our revision policy: We would like to draw your attention to changes in our revision policy that we have made in response to COVID-19 (https://elifesciences.org/articles/57162). Specifically, we are asking editors to accept without delay manuscripts, like yours, that they judge can stand as eLife papers without additional data, even if they feel that they would make the manuscript stronger. Thus the revisions requested above only address clarity and presentation.

----------

Information to help as you revise your submission:

* We will look forward to hearing from you with a revised article with tracked changes, and a response letter (uploaded as an editable file) describing the changes made in response to the decision and review comments.

* If source manuscript files have not already been provided (including figures in an RGB color space, uploaded individually), we will need them at the revision stage; further details are here: https://submit.elifesciences.org/html/elife_author_instructions.html#revised

* If your study includes bench research, please include a Key Resources Table within your resubmission. This is designed to highlight genetically modified organisms and strains, cell lines, reagents, and software that are essential to reproduce the results presented (but it is not designed to be a comprehensive list of all the materials and resources used).You should download the following template, which includes notes on completion as well as an example table: [https://elife-cdn.s3.amazonaws.com/author-guide/elifeKRTtemplatev2.docx] For relevant submissions, the completed table should be incorporated within your article file at the very beginning of the Materials and Methods section (examples in published articles are available at [https://elifesciences.org/articles/44199#s4] and [https://elifesciences.org/articles/45342#s4]).

* If your work involved the use of cell lines, please indicate in the Materials and Methods section of your manuscript if their identity has been authenticated, state the authentication method (such as STR profiling), and report the mycoplasma contamination testing status. Please consult our Journal Policies for further details: https://submit.elifesciences.org/html/elife_author_instructions.html#policies

* Advice about statistical reporting: in your revised paper, please ensure that any statistical analysis methods have been described and justified; ensure that raw data are presented in figures whenever informative to do so (typically when N per group is less than 10); for each experiment, ensure that you have identified the statistical tests used, the exact values of N, definitions of center, methods of multiple test correction, and dispersion and precision measures; and please report exact p-values wherever possible alongside the summary statistics and 95% confidence intervals (for all key questions and not only when the p-value is less than 0.05).

* If your revised article is accepted for publication, please note that authors can choose to have their accepted PDF published by eLife within a few days of acceptance, in advance of the version of record that will be subject to technical processing, editing, and author proofing. Please consider whether this is suitable for your work, so you can select the appropriate option at the time of resubmission. In the event that the study has potentially broad public relevance, we recommend you consult your press officer before making this decision.

----------

Please use the following link when you are ready to resubmit:

[https://submit.elifesciences.org/cgi-bin/main.plex?el=A5Hz3rZM1A1Bebn2I6A9ftdZzSJCsHv3LpEV9xfJzZeDwZ]

Best wishes,

Alan Talevi
Reviewing Editor, eLife

Amy Andreotti
Senior Editor, eLife

-----
eLife Sciences Publications, Ltd is a limited liability non-profit non-stock corporation incorporated in the State of Delaware, USA, with company number 5030732, and is registered in the UK with company number FC030576 and branch number BR015634 at the address Westbrook Centre, Milton Road, Cambridge, CB4 1YG, UK.
-----
